# How Ikaros and Aiolos homo- and heterodimers drive gene expression to ensure B cell development

**DOI:** 10.1101/2025.06.09.658591

**Authors:** Marie-Céline Deau, Beate Heizmann, Adina Aukenova, Quentin Heydt, Patricia Marchal, Céline Charvet, Stéphanie le Gras, Isabelle M.L. Billas, Dominic van Essen, Simona Saccani, Philippe Kastner, Susan Chan

**Author notes:** Correspondence: Susan Chan and Philippe Kastner. equal contribution (Marie-Céline Deau, Beate Heizmann). equal contribution (Adina Aukenova, Quentin Heydt). senior authors (Dominic van Essen, Simona Saccani, Philippe Kastner, Susan Chan).

## Abstract

Ikaros family transcription factors (TF) are major regulators of cell differentiation and function. They function as homo- and heterodimers, but the contribution of specific dimers to gene regulation is unknown. We investigated the function of Ikaros-Ikaros, Ikaros-Aiolos and Aiolos-Aiolos in B cell development, and show that the dimers are not equivalent. Ikaros homodimers are required in proB and large preB cells where they bind and regulate the expression of a surprisingly small number of genes related to cell adhesion and migration, predominantly as transcriptional repressors. In contrast, Ikaros-Aiolos heterodimers and Aiolos homodimers are required in small preB and immature IgM+ B cells to regulate 10-fold larger target gene repertoires involved in B cell receptor signaling and immune functions. Aiolos-containing dimers act mainly as repressors in small preB cells but activators in immature B cells, where Aiolos-Aiolos antagonizes Ikaros-Ikaros. Mechanistically, Aiolos binds more GGAA motifs with different flanking nucleotides than Ikaros, which depends on a single amino acid difference in zinc finger 3 of its DNA binding domain. These results illustrate how homo- and heterodimerization of homologous TF proteins can markedly impact binding specificity, target gene response and pathway modulation in cells of the same lineage.

## INTRODUCTION

The ability of transcription factors (TFs) to bind and function at target sites is a hallmark of gene regulation. Numerous TFs work as dimers, which allow them to double the length of their target sequence, increase DNA affinity and/or fine-tune specificity ^1^. Most studies to date have focused on the rules of dimerization and the structural compatibility between partners. How and whether dimerization imparts specificity in gene regulation remains unclear, mainly due to the difficulty in distinguishing different dimers at DNA.

The Ikaros family of TFs comprises four zinc finger (ZF) proteins whose genes probably arose from duplication events in a jawed vertebrate ancestor ^2,3^. Ikaros (encoded by *Ikzf1*) is highly homologous with Aiolos (*Ikzf3*), and Helios (*Ikzf2*) with Eos (*Ikzf4*). Most were discovered in the hematopoietic system where they are critical for cell differentiation and function, but have now been found to be essential for the development of multiple tissues ^4–6^. Ikaros family proteins are characterized by two C2H2 ZF DNA binding and dimerization domains ^7–9^, and act as homotypic dimers. It is currently unknown if the different homo- and heterodimers recognize the same target sequences, function similarly as activators or repressors, or regulate the same biological pathways.

Ikaros and Aiolos are differentially expressed in developing B lymphocytes ^10–12^. Ikaros is expressed throughout the B cell lineage from the earliest pre-proB cell stage in the bone marrow (BM), while Aiolos is detected preB and immature B cells. Accordingly, Ikaros is absolutely required for early B cell differentiation, by promoting B cell receptor (BCR) heavy and light chain rearrangements and the transition from large to small preB cells ^11,13–15^, and has been associated with gene repression ^12,16–18^. In contrast, the role of Aiolos in BM B cells is less clear. Aiolos knock- out (KO) B cells are phenotypically normal in the BM, but later show signs of autoimmune activation ^19–21^. Aiolos is also needed for the downregulation of the preBCR surrogate light chain λ5 in preB cells, indicating a unique role for which Ikaros is not sufficient ^22,23^. That Aiolos might heterodimerize with Ikaros in preB cells was recently suggested by the finding that a human *IKZF3* dominant-negative mutation, which negatively affects DNA binding, mimics the murine *Ikzf1* loss-of-function (LOF) phenotype, implying combinatorial control ^24^.

Unsurprisingly, Ikaros and Aiolos are implicated in multiple human pathologies including immunodeficiencies, autoimmunity and cancer. Germline *IKZF1* and *IKZF3* LOF mutations are associated with combined immunodeficiency (CID) and common variable immune deficiency (CVID) ^25–28^, and *IKZF1* variants have been found in patients with autoimmune disorders ^29^. *IKZF1* LOF mutations and deletions are prevalent in B cell precursor-acute lymphoblastic leukemias (BCP-ALL), highlighting the role of Ikaros as a tumor suppressor in early stage B cells ^30^. In contrast, gain-of-function (GOF) *IKZF3* alterations are linked to chronic lymphocytic leukemias and some B cell lymphomas ^31,32^, and overexpression of WT Ikaros and Aiolos promote cancer cell growth and survival in multiple myelomas ^33^.

The sequential appearance and non-redundant phenotypes of Ikaros family proteins in B cell differentiation suggested that selective binding and activities of individual dimers could play critical roles in regulating gene expression. In this study, we engineered a novel split epitope-based approach to directly address the function of Ikaros and Aiolos homo- and heterodimers in early B cells. We show that the different dimers are not equivalent, and that each is uniquely required at key transitional steps of B cell development.

## RESULTS

### Monitoring homo- and heterodimerization via fluorescence and epitope complementation

To study Ikaros and Aiolos dimers in early B cells, we first evaluated the presence of these TFs during differentiation, using the Hardy nomenclature to identify the different B cell stages ^34^. Ikaros proteins were easily detected in all BM B cells from Fractions A to F by flow cytometry (Fig. S1), in line with our previous results ^11^. In contrast, Aiolos appeared progressively: it was absent in Fraction A (pre/proB), present at low but increasing levels in Fractions B (early proB), C/C’ (late proB/large preB) and D (small preB), and robustly detected in Fractions E (immature IgM+ B) and F (mature recirculating IgM+B220++ B). These results suggested that all of the homo- and heterodimers exist starting at the large preB cell stage.

To monitor dimerization, we took advantage of a bimolecular fluorescence complementation (BiFC) approach, where Ikaros and Aiolos were fused at the C-terminus to the non-fluorescent V1 or V2 fragments of the Venus protein. Dimerization between the tagged proteins leads to the interaction of V1 and V2, and assembly into a fluorescent Venus reporter (Figs. S2, a and b). Importantly, we generated a novel monoclonal antibody (Mab199) that recognizes an epitope present on the assembled (and native) Venus, but not the V1 or V2 fragments (DvE and SS, in preparation, and see below); Mab199 is suitable for immunoprecipitation and ChIP-sequencing.

We then established an inducible expression system in the murine BH1 preB cell line ^11^. BH1 cells were derived from short-term cultures of Ikaros KO BM cells; they are dependent on IL7 for growth, blocked at the large preB cell stage, and their differentiation can be rescued upon Ikaros re-expression. Here, BH1 cells were further engineered to express combinations of tagged Ikaros and Aiolos under doxycycline (dox) control (Fig. S2b). The stable BH1-derived cell lines, expressing Ikaros-V1 and Ikaros-V2 (hereafter called "Ikaros-Ikaros"), Aiolos-V1 and Aiolos-V2 ("Aiolos-Aiolos"), Ikaros-V1 and Aiolos-V2 ("Ikaros-Aiolos"), or only the tetracycline-dependent reverse transactivator (rtTA), were retrovirally produced and subjected to antibiotics selection. Upon dox treatment, cells from the Ikaros-Ikaros lines were predicted to express the fluorescent V1-V2 (as well as the non-fluorescent V1-V1 and V2-V2) Ikaros homodimers. Similarly, cells from the Aiolos-Aiolos lines were predicted to express the fluorescent V1-V2 (as well as the non-fluorescent V1-V1 and V2-V2) Aiolos homodimers. Lastly, cells from the Ikaros-Aiolos lines were predicted to express the fluorescent Ikaros-V1-Aiolos-V2 heterodimers, and the non-fluorescent Ikaros-V1-Ikaros-V1 and Aiolos V2-Aiolos-V2 homodimers. Only the Venus+ dimers can be detected by Mab199.

Dox treatment induced Venus expression in >85% of the cells in every line after 16h (Fig. S2, c and d), indicating successful V1-V2 dimerization. Venus+ Ikaros-Ikaros dimers accumulated at centromeric heterochromatin, like untagged Ikaros (Figs. S2d and S3a) ^35^, suggesting that the Venus reporter did not inhibit Ikaros localization in the nucleus. Ikaros-Ikaros preB cells underwent differentiation after 5 days of dox treatment (Fig. S3b), as measured by the appearance of surface B cell receptors (BCR), similar to our previous system with tamoxifen-induced Ikaros ^11^, demonstrating that the tagged TFs can efficiently regulate multiple gene pathways. Further, Venus+ Ikaros homodimers bound the Ikaros target region at the *Cish* gene ^36^, in ChIP experiments using Mab199 or a control anti-Ikaros antibody (Fig. S3c), indicating specific DNA binding. These data showed that the V1- and V2-tagged Ikaros TFs were functional. To determine if the V1-V2 interaction itself could lead to artifactual dimerization, we expressed WT Ikaros-V1 and Ikaros-V2, or mutant IkΔDD-V1 and IkΔDD-V2 lacking the Ikaros dimerization domain, in 293T cells and immunoprecipitated the proteins with Mab199 (Fig. S3d). Only WT Ikaros-Ikaros was detected, and not IkΔDD-IkΔDD, indicating that the V1-V2 interaction required TF dimerization. We also asked if IkΔDD-V1 and IkΔDD-V2 could bind the synthetic BS4 DNA probe containing two high affinity Ikaros binding motifs ^7^ as a dimer, by EMSA (Fig. S3e). Only WT Ikaros-Ikaros bound BS4 as a dimer, which could be supershifted with Mab199. To determine if the V1-V2 interaction increased dimer stability on DNA, we tested the ability of excess unlabeled BS4 to displace Ikaros-Ikaros binding (Fig. S3f); TF binding was similarly displaced between untagged and Venus-tagged Ikaros samples. Thus, the V1-V2 complementation alone did not induce dimerization or stabilize Ikaros dimers on DNA, though we could not exclude that it may help to maintain the dimer state once dimerization occurred.

To normalize Venus expression across cell lines, and express Ikaros and Aiolos at physiological levels, we tested different protocols of dox treatment (Fig. S4, a and b). "High dox" (2 μg/ml) induced similarly high Venus levels and TF overexpression. "Low dox" (50 ng/ml for Ikaros-Ikaros, 35 ng/ml for Ikaros-Aiolos and Aiolos-Aiolos) induced lower Venus levels and physiological TF expression, using primary cells where these TFs were easily detectable [BM immature B cells (CD19+ IgM+) to standardize Ikaros, splenic CD19+ B cells to standardize Aiolos]. Thus, low dox treatment was used for the majority of the assays below.

### Ikaros homodimers bind a small subset of the targets recognized by Aiolos

We analyzed the binding of the Ikaros and Aiolos homo- and heterodimers to the chromatin of the modified BH1 cells cultured in low or high dox for 16h, by ChIP-sequencing using Mab199. As controls, the binding profiles of the Venus+ dimers were compared with those of Ikaros from cells expressing Ikaros V2 (IP-ed with anti-Ikaros), and Aiolos from cells expressing Aiolos-V2 (IP-ed with anti-Aiolos). Similar results were obtained between Mab199 and anti-Ikaros or anti-Aiolos (Fig. S5, a-d), indicating that the Venus+ dimers recognized bona fide targets.

Unexpectedly, principal component analysis revealed a clear segregation in DNA binding amongst the different dimer pairs regardless of dox treatment (Fig. 1a), which indicated specific motif recognition that was not markedly altered or enhanced by TF overexpression (Fig. S5e). Ikaros-Ikaros bound far fewer targets than Ikaros-Aiolos or Aiolos-Aiolos (Fig. 1b), and the median height of the Ikaros-Ikaros peaks was lower than those of the other dimers (Fig. 1c). Nonetheless, visual scanning of the genome browser tracks revealed that the Ikaros-Ikaros peaks were not uniformly low, and ranged from comparatively high when compared with the Aiolos-containing dimer peaks (Fig. S5a, black arrowheads), to almost non-existent (white arrowheads). In addition, reduced Ikaros-Ikaros binding was observed in both low and high dox-treated cells, where the mean fluorescence intensity of Venus in the high dox-treated Ikaros-Ikaros cells was higher than that of the low dox-treated Aiolos-Aiolos cells (see Fig. S4a), indicating sufficient TF levels.

**Figure 1.**
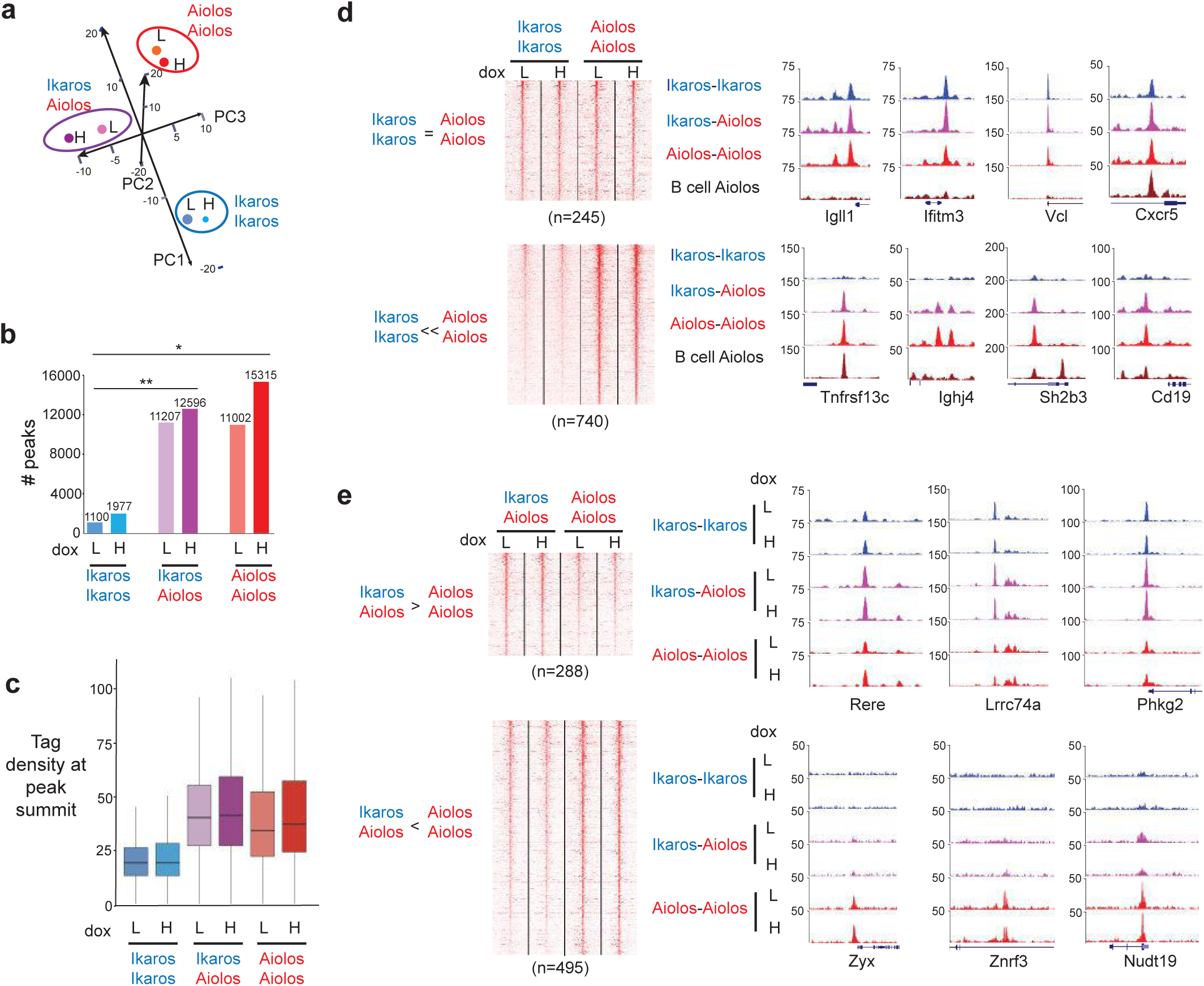
Differential binding by Ikaros and Aiolos homo- and heterodimers. **(a)** PCA of the homo- and heterodimer peaks in cells cultured for 24 h with low or high dox, followed by ChIP-seq with Mab199 anti-Venus. L: low dox; H: high dox. **(b)** Number of peaks detected by MACS2 in the cells expressing the indicated tagged dimers. *p<0.05; **p<0.01 (Student’s t-test). **(c)** Box plots showing the distribution of the tag densities at peak summits (determined from 15x10^6^ reads in each sample) out of 701 common peaks detected in the 6 ChIP-seq samples. **(d)** Illustration of peaks commonly bound by Ikaros and Aiolos homodimers (top), or selectively bound by Aiolos-containing dimers (bottom), as detected by Mab199. Peaks shown in the heatmaps were selected as: commonly bound peaks, where pileup Ikaros >30, I Log2 FC I (Ikaros vs Aiolos)<0.2; adj. p>0.2 (DEseq2); Aiolos-biased peaks, where pileup Aiolos>30, Log2 FC (Ikaros vs Aiolos)<-1.5; adj. p<0.05. UCSC genome browser tracks correspond to high dox samples, generated from 15x10^6^ reads. Control tracks show Aiolos binding in splenic CD19+ B cells, as detected by the anti-Aiolos Ab. Scales correspond to tag numbers. Gene names are indicated below the tracks. **(e)** Illustration of peaks differentially bound by Ikaros-Aiolos vs. Aiolos-Aiolos, as detected by Mab199. Heatmaps correspond to peaks identified as differential by DEseq2 (I Log2 FC I >0; adj. p<0.05).

Of the 17,911 targets identified by MACS2 (low and high dox samples were counted as duplicates), 10,535 had higher Aiolos-Aiolos peaks compared with Ikaros-Ikaros according to DEseq2 (Log2 FC>0.5; adj. p<0.05) (not shown), 245 had peaks of similar height with both homodimers (Fig. 1d, top), and only 46 had higher Ikaros-Ikaros peaks (Fig. 1e, see *Phkg2*, and not shown). Further, 740 targets were bound by Aiolos-Aiolos (Aiolos-biased) and not Ikaros- Ikaros; these included *Tnfrsf13c* (encoding the BAFF receptor), *Ighj4*, *Sh2b3* (encoding the LNK adapter), *Cd19*, *Tlr9*, *Cd79a* and *Blnk* (Figs. 1d, bottom, and S5f). Ikaros-Ikaros only peaks were never detected. When Ikaros-Aiolos peaks were compared with Aiolos-Aiolos ones, >99% were present at the same sites, though the heterodimer peaks were higher at 288 sites and lower at 495 (Fig. 1e). Importantly, similar conclusions were deduced from the results of ChIP-seq analyses using anti-Ikaros vs. anti-Aiolos antibodies (see Fig. S5, a-d), though a direct comparison was not possible due to the use of different antibodies. These results showed that >2700 of the targets bound by Ikaros-Aiolos or Aiolos-Aiolos in BH1 cells were bound by endogenous Aiolos in splenic CD19+ B cells (Figs. 1d and S5, f and g), where both Ikaros and Aiolos are highly expressed, suggesting that Aiolos binds many of the same targets in preB and mature B cells.

Thus, Ikaros and Aiolos have different DNA binding specificities: the Ikaros homodimer binds a low number of motifs, while the heterodimer and the Aiolos homodimer recognize ∼10- fold more target sequences.

### Aiolos evolved to recognize an expanded range of target sequences

To understand why Aiolos bound more targets than Ikaros, we analyzed the sites selectively bound by Aiolos, which frequently contain 1-5 conserved GGAA motifs under the peak summit (Fig. S6a, and not shown), and compared the binding of Ikaros vs. Aiolos to three Aiolos-biased sequences from the *Tnfrsf13c*, *Ighj4* and *Sh2b3* loci by EMSA, using nuclear extracts of COS cells expressing full-length Ikaros or Aiolos (Fig. 2a). BS4 and a modified probe from the *Cish* gene ^36^, both of which are bound strongly by Ikaros and Aiolos, were used as controls. These experiments showed that Aiolos bound the *Tnfrsf13c*, *Ighj4* and *Sh2b3* probes better than Ikaros even when using linear DNA fragments.

**Figure 2.**
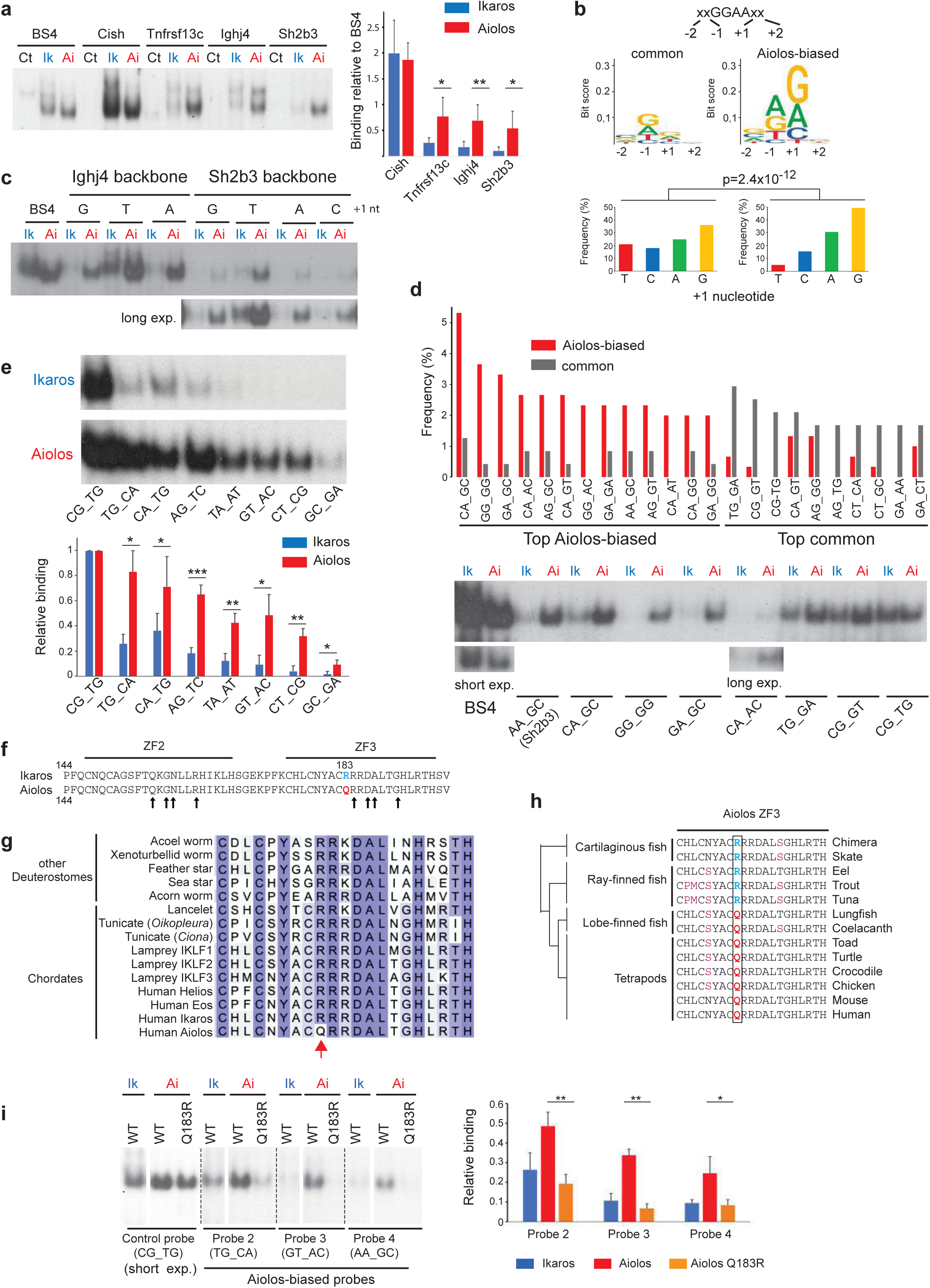
Distinct motifs recognized by Ikaros vs. Aiolos. **(a)** Left: Representative EMSA showing binding of Ikaros and Aiolos to control probes (BS4, *Cish*) and probes derived from the *Tnfrsf13c*, *Ighj4* and *Sh2b3* loci (see Fig. S6a). Control (Ct) samples correspond to nuclear extracts from Cos cells transfected with the empty pTL2 expression vector. Right: Comparison of relative binding of Ikaros and Aiolos on the indicated probes. Values represent binding relative to that on the BS4 probe. Data are from 3 experiments. *p<0.05; **p<0.01 (Student’s t-test). **(b)** Top: Nucleotide enrichment among the 4 nts adjacent to the GGAA core motifs found under the Aiolos-specific peaks or under peaks commonly bound by Ikaros and Aiolos (see Fig. S6d) (motifs generated with ggseqLogo). Bottom: Frequency of each base at the +1 nt. P-value was calculated with the Chi-square test, using the distributions seen with the common Ikaros and Aiolos peaks as reference. **(c)** Influence of the +1 nt on Ikaros and Aiolos binding. Binding was tested by EMSA to probes derived from the *Ighj4* or *Sh2b3* backbones in which the +1 nts were converted to the indicated nucleotides (2 motifs for *Ighj4*, 1 motif for Sh2b3). Note that the "C" conversion is not shown for the *Ighj4* probe because it led to strong non-specific binding by a nuclear factor present in Cos cells. Long exp: long exposure. **(d)** Top: Frequency of the most frequent 4-nt environments surrounding the GGAA motifs from Aiolos-biased targets vs. common targets (see Fig. S6d for motif selection). Bottom: Impact of the top nt contexts from Aiolos-biased and common targets on TF binding to *Sh2b3*-derived probes, by EMSA. **(e)** Binding of Ikaros and Aiolos to *Sh2b3*-derived probes with various nucleotide contexts. The 4-nt contexts of the probes (excluding the control probe) were chosen to overlap at most by one nt in any pairwise comparison. Top: Representative EMSA. Bottom: Comparison of relative binding from 3 independent experiments. Data were relative to binding observed with the probe containing the CG_TG context, which binds Ikaros and Aiolos similarly. *p<0.05; **p<0.01; ***p<0.001 (Student’s t-test). **(f)** Alignment of the DNA binding domain ZF2 and ZF3 of murine Ikaros and Aiolos. Arrows point to the residues predicted to interact with DNA. **(g)** Alignment of Ikaros-related ZF3 sequences from the indicated species. **(h)** Alignment of Aiolos ZF3 in representative vertebrate species. For (g) and (h), see ^3^ for references of the sequences used for the alignments. **(i)** Left: Representative EMSA comparing binding of full-length WT Ikaros and Aiolos vs. Aiolos Q183R. Probes used were taken from (e). Note that the image came from one EMSA experiment, but some irrelevant lanes were cropped out from the autoradiogram in this panel. Right: Comparison of relative binding from 3 independent experiments. Binding was normalized to that of the control probe CG_TG. *p<0.05; **p<0.01; ***p<0.001 (Student’s t test).

We next addressed the importance of the flanking sequences. The *Sh2b3* probe was chosen as the model system because it contained only one GGAA motif recognized by one protein subunit, making it simpler to manipulate. Nevertheless, it was bound by Aiolos in dimer form, as shown with extracts containing Aiolos-V1 and Aiolos-V2, and Mab199 to supershift the V1-V2 complex (Fig. S6b, and manuscript in preparation). The *Sh2b3* probe was mutated 2 nts up- and downstream of the GGAA motif to match those of the BS4 or *Cish* motifs, and subjected to EMSA with extracts containing full-length Ikaros or Aiolos (Fig. S6c). These changes allowed Ikaros to bind to the modified *Sh2b3*, but also increased Aiolos binding, indicating that the flanking sequences contributed to Ikaros and Aiolos specificity and overall strength of binding. We then analyzed 84 common and 132 Aiolos-biased peaks in situ, by studying the GGAA motifs within 90 bps of the peak summits, which revealed 238 and 301 motifs, respectively (Fig. S6d). Similar nucleotide usage was observed at nucleotides -2, -1 and +2 for both types of motifs (Fig. S6e), but a strong bias was seen against a T at +1 among the Aiolos-biased motifs (Fig. 2b). To test the importance of this pyrimidine, we mutated the +1 nucleotide to a T in the *Sh2b3* and *Ighj4* probes, and tested the consequences by EMSA (Fig. 2c). Contrary to expectations, a T at +1 did not reduce Aiolos binding but instead enhanced the binding of both Ikaros and Aiolos, indicating that Aiolos specificity was due to the selection of lower affinity sequences to which Ikaros cannot bind. Indeed, Aiolos was generally less stringent than Ikaros for binding to a variety of DNA fragments, as the top Aiolos-biased motifs had flanking sequences that were recognized selectively by Aiolos in EMSAs (Fig. 2d), while all of the top common motifs were bound by both Ikaros and Aiolos. Aiolos also bound better than Ikaros by 2- to 5-fold to *Sh2b3*-modified probes containing randomly chosen flanking nucleotides that minimally overlapped (Fig. 2e). Interestingly, sequence-specific deformability analysis suggested that the flanking nucleotides from probes that bound both Ikaros and Aiolos had more flexible nucleotide steps (in particular Y-R steps in the 5’ region) than those from Aiolos-biased probes, as evidenced by the roll angle values (Fig. S7a).

To determine if the Ikaros-Aiolos heterodimer had its own specificity, we tested the binding of V1- and V2-tagged Ikaros-Ikaros, Aiolos-Aiolos and Ikaros-Aiolos to probes that were differentially bound by Ikaros and Aiolos, and used Mab199 to supershift (Fig. S6f). Ikaros-Aiolos bound the TG_CA and AG_TC probes as efficiently as Aiolos-Aiolos, but was less efficient with TA_AT and GT_AC, indicating that Ikaros-Aiolos binding was influenced by both the Ikaros and Aiolos subunits.

To investigate the protein sequence differences between Ikaros and Aiolos, we focused on the amino acids in ZF2 and -3 of the DNA binding domain ^7^. These zinc fingers possess only one amino acid difference at aa183 in ZF3 between Ikaros (R) and Aiolos (Q) (Fig. 2f), which is immediately N-terminal of a critical arginine (aa184) required for base-specific DNA contact ^37^. Interestingly, R183 is present in the other Ikaros family members (Helios, Eos), and in Ikaros homologs from all deuterostome classes (Fig. 2g). It is also the ancestral residue of Aiolos, as R183 is found in Aiolos from cartilaginous and ray-finned fish, but not from fish related to tetrapods (lungfish, coelacanth) or other tetrapods, which have a glutamine at this position (Fig. 2h), indicating that Aiolos Q183 evolved from a highly conserved arginine at the emergence of tetrapods and was subsequently conserved. To determine if Q183 was important for DNA binding by Aiolos, we mutated this residue in Aiolos (Q183R), and compared the binding of WT and mutant Aiolos to the Aiolos-selective *Sh2b3* sequence (AA_GC) and its derivatives (TG_CA, GT_AC). Strikingly the Q183R mutation abolished Aiolos specificity to all 3 probes (Fig. 2i), demonstrating that Q183 is required for the extended binding capacity of Aiolos. Structural modeling indicated that Ikaros R183 likely forms electrostatic interactions with the DNA phosphate backbone, while Aiolos Q183 appears to interact intramolecularly with the protein itself (Fig. S7b). Consistent with the sequence-specific deformability analysis (see Fig. S7a), these differences imply that Ikaros requires DNA to be bendable for optimal binding, while effective Aiolos binding may rely less on DNA shape. This could explain why Ikaros target sequences tend to be intrinsically more flexible than those recognized by Aiolos.

Collectively, these results demonstrated that Aiolos binds more GGAA motifs with different flanking nucleotides than Ikaros, which depends on a single amino acid difference in ZF3 that evolved in vertebrates.

### Ikaros is associated with gene repression and Aiolos with activation

To study target gene regulation, we evaluated the transcriptomes of BH1 cells expressing Ikaros- Ikaros, Ikaros-Aiolos, Aiolos-Aiolos, or rtTA after 24h of low dox, by RNA-sequencing. In the cells expressing Ikaros-Aiolos, all combinations of dimers should co-exist, which made it harder to pinpoint heterodimer function but also mimics the physiological state of small preB cells. The differentially expressed genes (I Log2 FC >0.5 I; adj. p<0.05) were further filtered to reveal only the target genes bound by at least one of the dimers (hereafter named DETGs).

The expression of Ikaros-Ikaros or Aiolos-Aiolos impacted 1029 or 2051 DETGs, respectively (Figs. 3a), with an intermediate response to Ikaros-Aiolos. In addition, Ikaros-Ikaros repressed more genes than it activated, and vice versa for Aiolos-Aiolos. Scatter plot analysis showed that 230/284 (>80%) of the commonly upregulated DETGs were more activated by Aiolos-Aiolos (Fig. 3b), while 260/412 (>63%) of the commonly downregulated DETGs were more repressed by Ikaros-Ikaros. Even among the antagonistically regulated DETGs, Aiolos-Aiolos activated more genes repressed by Ikaros-Ikaros than the other way around (15 vs. 2, respectively).

**Figure 3.**
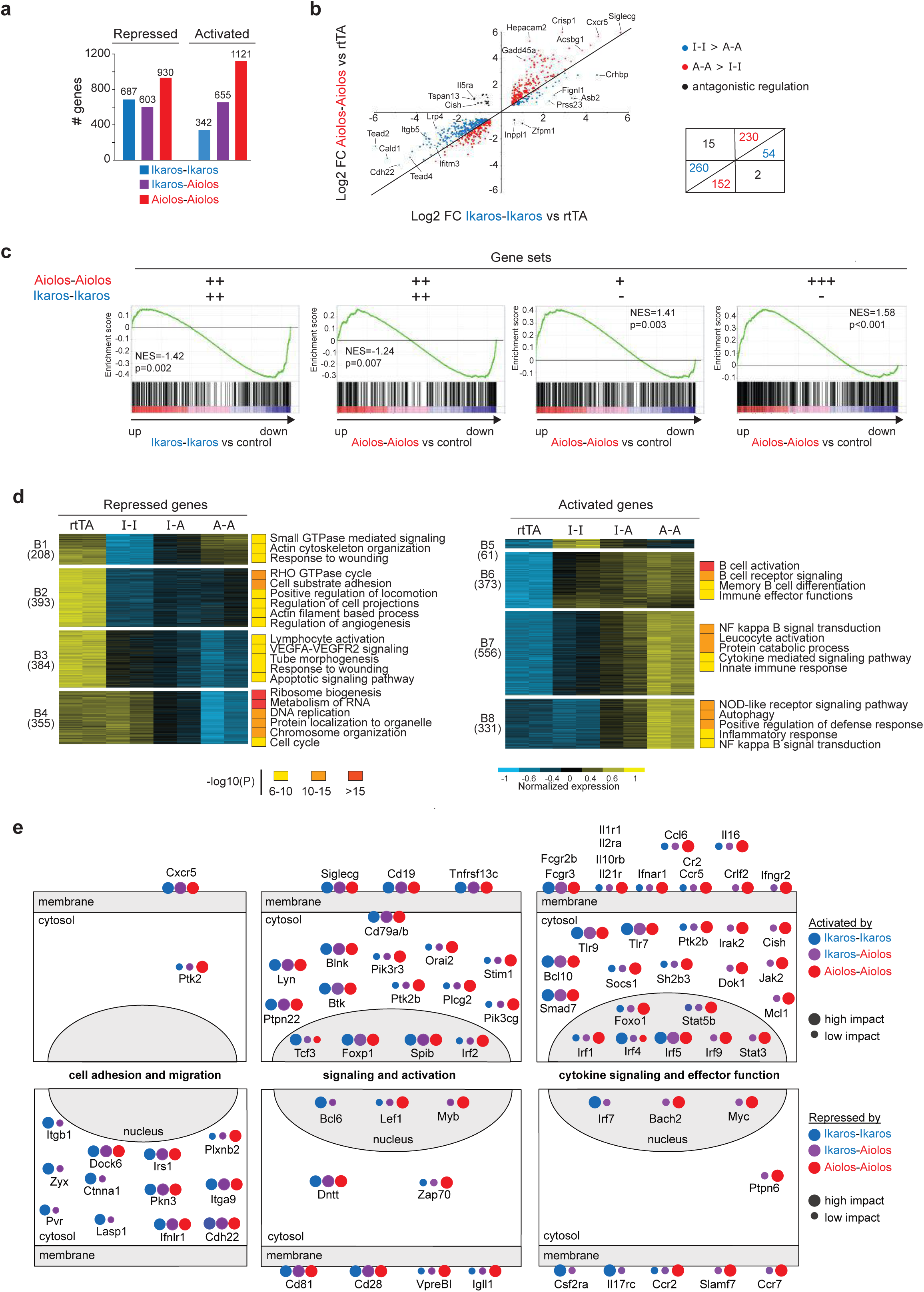
Ikaros and Aiolos differentially regulate gene expression in BH1 cells. **(a)** Number of directly activated and repressed genes after dox induction (24 h, low dox) of the Ikaros-Ikaros, Ikaros-Aiolos, and Aiolos-Aiolos cell lines. The analysis was restricted to genes bound by any dimer. Selection criteria: I Log2 FC >0.5 I; adj. p<0.05. **(b)** Scatter plot showing the Log2 FC (with respect to the rtTA samples) of the genes that were regulated by both Ikaros and Aiolos (I Log2FC I >0.5; adj. p<0.05) in the Ikaros-Ikaros and Aiolos-Aiolos cells. Red, blue and black dots respectively represent genes regulated more strongly by Aiolos, Ikaros, or antagonistically by both factors. **(c)** GSEA of gene sets corresponding to distinct groups of Ikaros-Ikaros or Aiolos-Aiolos bound genes (by ChIP-seq), and ranked Aiolos-regulated or Ikaros-regulated genes. +++: Aiolos-Aiolos pile-up values >70 in both Aiolos-Aiolos (HD and LD) ChIP-seq samples; ++: pile-up values >25 in all Ikaros-Ikaros and Aiolos-Aiolos ChIP-seq samples; +: pile-up values <30 in both Aiolos-Aiolos samples; -: peak not detected in both Ikaros-Ikaros samples (see Fig. S8a for representative peaks in the selected groups). 500 randomly selected genes from each group were used for the GSEA analysis. NES: Net enrichment score; The p-value corresponds to the proportion of enrichments with a higher enrichment score obtained with 1000 random permutations of the ranked list. **(d)** Heatmap of the relative expression of the genes regulated by Ikaros (I-I), Aiolos (A-A) or their combination (I-A) (I Log2 FC I >0.5; adj. p<0.05 by K-means clustering). Numbers of genes in each cluster are indicated. The top pathways enriched in each cluster are indicated (Metascape; except for cluster B1 for which no clear pathway emerged). **(e)** Representation of genes (placed according to the location of the encoded protein) and their regulation by Ikaros-Ikaros, Ikaros-Aiolos or Aiolos-Aiolos of representative genes from major pathways regulated by Ikaros and/or Aiolos.

We asked how the targets commonly bound by Ikaros and Aiolos were regulated compared with those selectively bound by Aiolos (low and high binding peaks) (Fig. S8a), using Gene Set Enrichment Analyses (GSEA) (Fig. 3c). This demonstrated that the commonly bound targets were preferentially repressed by Ikaros or Aiolos. On the other hand, the Aiolos-biased targets were mostly activated by Aiolos, regardless of binding affinity. The DETGs were further studied by K-means clustering and grouped into 8 similarly regulated gene clusters (Fig. 3d). Only 2 clusters (B2, B6, together corresponding to 29% of the DETGs) were similarly regulated by Ikaros-Ikaros or Aiolos-Aiolos (Fig. S8b, see *Ifitm3* and *Tnfrsf13c*). The other clusters showed specific repression or activation by either Ikaros-Ikaros or Aiolos-Aiolos (Fig. S8b, see *Lrp1*, *Myc*, *Myl4*, *Ccl9*). Some DETGs were antagonistically regulated (Fig. S8b, see *Cish*). DETGs with high Ikaros-Ikaros peaks were associated with Ikaros-mediated repression, while those with Aiolos- biased peaks were linked to Aiolos-mediated activation (Figs. S8, c and d). Conversely, the genes specifically activated by Ikaros (cluster B5) or repressed by Aiolos (B4) were respectively enriched for binding by Ikaros-Ikaros or Aiolos-containing dimers (Fig. S8d). These analyses indicated that Ikaros-Ikaros and Aiolos-Aiolos are functionally distinct, and that different binding patterns are associated with distinct gene regulation outcomes.

We next asked if the commonly or specifically regulated DETGs were associated with distinct biological processes. Indeed, the commonly regulated targets most strongly repressed by Ikaros-Ikaros (clusters B1 and B2) were enriched for processes like cell adhesion and locomotion (Fig. 3d). On the other hand, the genes specifically repressed by Aiolos-Aiolos (cluster B4) were enriched for intracellular and metabolic processes and the cell cycle; for instance, the key cell cycle regulator *Myc* was selectively repressed by the Aiolos homodimers (see Fig. S8b). Activated genes were mostly involved in immune effector functions like B cell activation, B cell receptor signaling, cytokine signaling, and inflammatory responses (Fig. 3d); they were strongly enriched among the genes preferentially induced by the Aiolos-containing dimers (B7 and B8), indicating a major role for Aiolos in activating the immune response. Figure 3e summarizes the top processes, and some of their associated genes, repressed or activated by the three dimer pairs. Altogether, our results indicated that Ikaros homodimer binding is associated with gene repression related to cell adhesion and cell movement, while heterodimer and Aiolos homodimer binding are linked to gene activation involved in immune effector functions. We next asked if these dimers functioned similarly in primary B cells.

### Sequentially appearing dimers regulate distinct gene expression programs in primary B cells

Given that Ikaros is ubiquitously expressed in all bone marrow B cells and Aiolos begins to be expressed in late proB/large preB cells, we predicted that Ikaros-Ikaros would be important in the early pre/proB to large preB cell stages (Fractions A-C/C’), while Ikaros-Aiolos and Aiolos-Aiolos would be important in the more differentiated small preB to immature B cell stages (Fractions D-E).

We first asked if the target genes in the BH1 cells were expressed in primary B cells at stages where the dimers were predicted to act. The transcriptomes of WT B cell fractions B, C/C’, D and E were evaluated by RNA-seq, and the genes differentially expressed (I Log2 FC >0.5 I; adj. p<0.05) between Fractions B and C/C’, C/C’ and D, or D and E were filtered to look at the BH1 DETGs from clusters B1-B8 (Figs. 4a and S9). This revealed a striking association between the BH1 targets repressed or activated by the different dimers and the genes down- or upregulated at the early and late B cell stages. Specifically, the BH1 DETGs repressed (or activated) by Ikaros-Ikaros were significantly enriched among the down- (or up-) regulated genes between B and C/C’, and those repressed (or activated) by Ikaros-Aiolos or Aiolos-Aiolos were down- (or up-) regulated between C/C’ and D, and/or D and E. Indeed, enrichment for the Aiolos- regulated genes was particularly high among the genes differentially expressed between C/C’ and D, suggesting a major switch to Aiolos-containing dimer function at the small preB cell stage.

**Figure 4.**
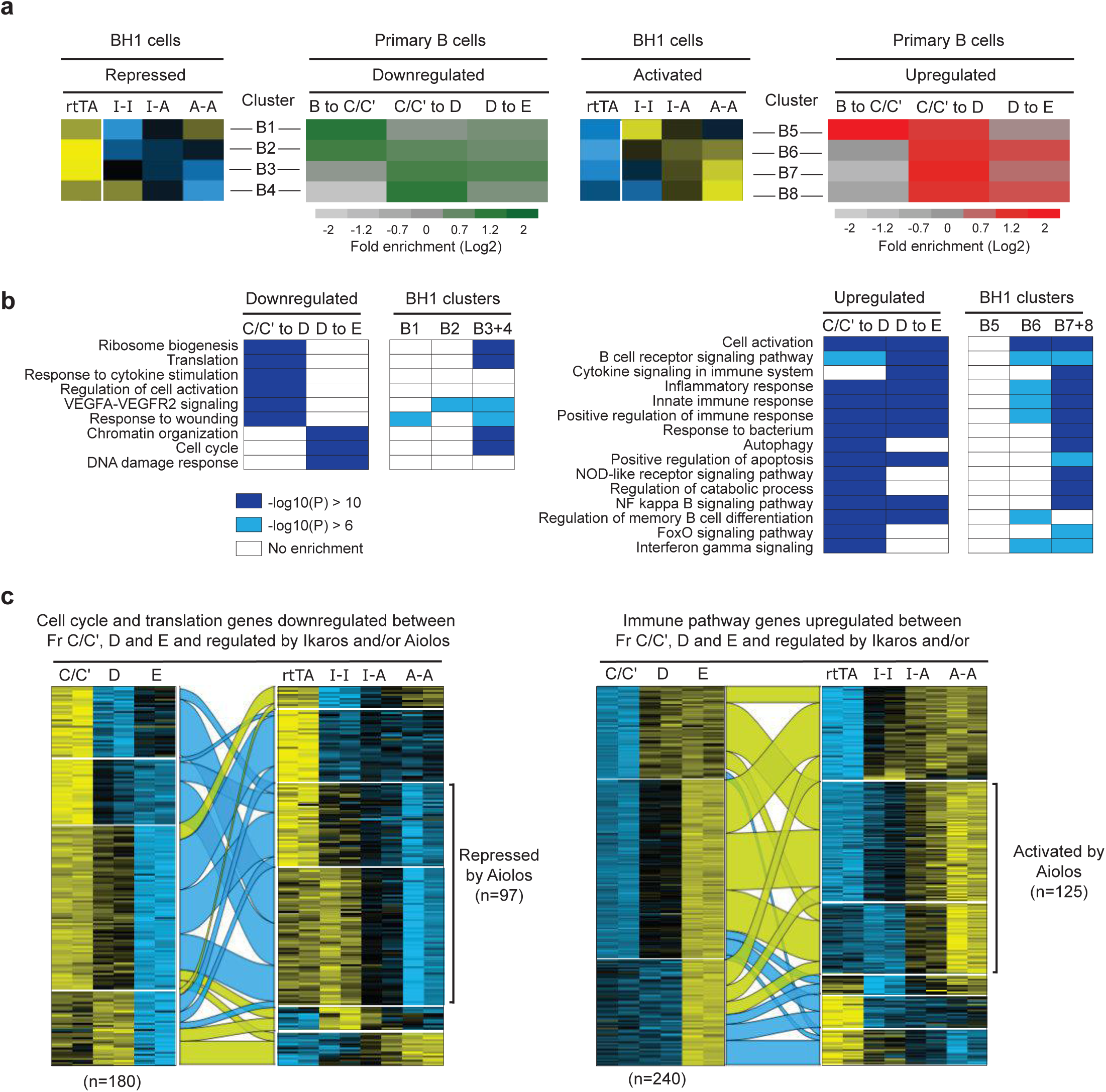
Ikaros and Aiolos regulate specific gene sets during BM B cell differentiation. **(a)** Enrichment of genes from the 8 gene clusters regulated by Ikaros or Aiolos in BH1 cells (see Fig. 3d) among genes that are up- or downregulated at the transitions between Fractions B and C/C’, C/C’ and D, or D and E. Genes upregulated during the developmental transitions between population X and Y were selected with the following criteria: I Log2 FC I >0.7; adj. p<0.05. Only the genes detected in both RNA-seq datasets were considered. Genes annotated as "Gene Model" or pseudogenes were excluded. See Fig. S9a for gene numbers and statistics. Colors for the BH1 clusters depicted on the left were calculated from the average normalized expression values of genes from the cluster. **(b)** Major pathways associated with genes induced or repressed between Fractions C/C’ and D, or D and E, and their enrichment in the indicated BH1 gene clusters. **(c)** Left: Heatmaps showing the expression in primary and BH1 cells of 180 genes related to cell cycle or translation-related pathways that were repressed between Fractions C/C’, D and E, and regulated by Ikaros and/or Aiolos in BH1 cells. Genes from the following pathways were considered: ribosome biogenesis, translation, Myc activation pathway, tRNA metabolic process, chromosome organization, cell cycle, chromosome segregation, DNA metabolic process, chromosome maintenance. Right: Heatmaps showing the expression in primary and BH1 cells of 240 genes related to immune-related pathways that were induced between Fraction C/C’, D and E, and regulated by Ikaros and/or Aiolos in BH1 cells. Genes from the following pathways were considered: cell activation, positive regulation of immune response, innate immune response, cytokine signaling in the immune system, autophagy, NFkB signal transduction pathway, response to bacterium, IFNγ signaling pathway, NOD-like receptor pathway, regulation of immune effector functions, inflammatory response, adaptive immune response.

We also determined if the differentially expressed genes in the primary cells were enriched for the Ikaros- or Aiolos-dependent pathways identified in BH1 cells, by assigning all of the genes differentially regulated between Fractions B to E to GO biological pathways, and comparing the top pathways with those from the BH1 clusters. For the B to C/C’ transition, this strategy was mostly unsuccessful, as the top pathways were mostly unrelated to those from the BH1 system. Nevertheless, of the 840 downregulated genes, 102 were repressed by Ikaros in clusters B1 and B2, and 31 of these (30%) belonged to pathways related to cell adhesion or motility (Fig. S9b), suggesting Ikaros-Ikaros dependent repression in large preB cells. A similar enrichment was not found among the ∼500 upregulated genes, where few genes were shared between primary and BH1 cells (10 belonged to cluster B5), and no pathways found.

Many more genes and pathways were affected by Ikaros and Aiolos between Fractions C/C’ and E. Of the 3568 down- and 3000 upregulated genes in the primary cells, 625 and 676 respectively were similarly regulated in the BH1 clusters. Indeed, the pathways associated with the C/C’ to D and D to E transitions resembled those of clusters B3+4 and B7+8 (Fig. 4b), which comprised the genes most affected by Ikaros-Aiolos and Aiolos-Aiolos. The pathways associated with gene downregulation were related to cell proliferation and growth (eg. ribosome biogenesis, translation, chromatin organization, cell cycle), and those associated with gene upregulation were related to effector immune cell responses (eg. cell activation, cytokine signaling, immune responses, autophagy, NF-kB signaling), further supporting the role of Aiolos-containing dimers at later steps of differentiation.

To determine how the genes from the top down and up pathways were regulated by the different dimer pairs in Fractions C/C’ to E, we focused on the genes involved in cell cycle/cell growth and immune functions (Fig. 4c). In all, 180 genes from the cell cycle/cell growth pathways were downregulated in Fractions D and E, and repressed by the dimers in BH1 cells. Conversely, 240 genes related to immune pathways were upregulated in D and E, and activated by the dimers in BH1 cells. These genes were then analyzed separately by K-means clustering, and the relationships between the clusters were plotted in alluvial flow diagrams. The results showed that about half of the genes down- or upregulated between C/C’ and D were those most sensitive to Ikaros-Aiolos and Aiolos-Aiolos in the BH1 cells, in line with the predicted appearance of these dimers at the small preB cell stage.

Thus, the genes differentially expressed at specific B cell developmental transitions reflect the regulatory specificities of the predicted Ikaros and Aiolos dimer types in those cellular stages. These findings suggested that the individual dimers, and particularly the Aiolos-containing dimers, were essential for the latter steps of B cell development.

### Ikaros-Aiolos and Aiolos homodimers are required for late B cell differentiation

To test the hypothesis that the Aiolos-containing dimers were required in small preB and immature B cells, we studied LOF mutants and focused on the transcriptomes of Fractions D and E cells. If the heterodimers were essential at these stages, then reducing either Ikaros or Aiolos should perturb the expression of their target genes. To reduce Ikaros, we took advantage of the Ikaros +/L mouse line in which one *Ikzf1* gene was replaced with the hypomorphic L allele; the +/L hematopoietic cells expressed Ikaros at ∼60% of WT levels ^38,39^. Though only the most strongly deregulated genes could be evaluated in this system, homozygous Ikaros L/L animals could not be studied because the L/L and KO strains display drastic B cell differentiation blocks at multiple stages ^13,38,40^. To knock out Aiolos, we generated a novel Aiolos KO (null) mouse line in which the last *Ikzf3* exon was deleted by CRISPR/Cas9-mediated mutation. The bone marrow B cell compartment of the Aiolos KO mice was similar to that of WT animals, with slight increases in the proportions of Fractions B and E, and a decrease in Fraction F, cells (Fig. S10a).

The transcriptomes of Fractions D and E cells from the WT, Ikaros +/L and Aiolos KO bone marrow were determined by RNA-seq. In these experiments, we used a lower FC threshold to select for the differentially expressed genes (| log2 FC | >0.2; adj. p<0.05) because many of the genes exhibited significant but modest changes. We then restricted our analysis to the BH1 DETGs (clusters B1-B8). These results showed that the Ikaros +/L mutation affected fewer genes than the Aiolos KO, as expected (Fig. 5a), though the pattern of deregulation was similar between Ikaros and Aiolos LOF in Fraction D where more genes were upregulated than down. Interestingly, this pattern was flipped in Fraction E, as the differentially expressed genes were mostly upregulated in the Ikaros +/L samples (68 up vs. 11 down), but prominently downregulated in the Aiolos KO (326 up vs. 490 down).

**Figure 5.**
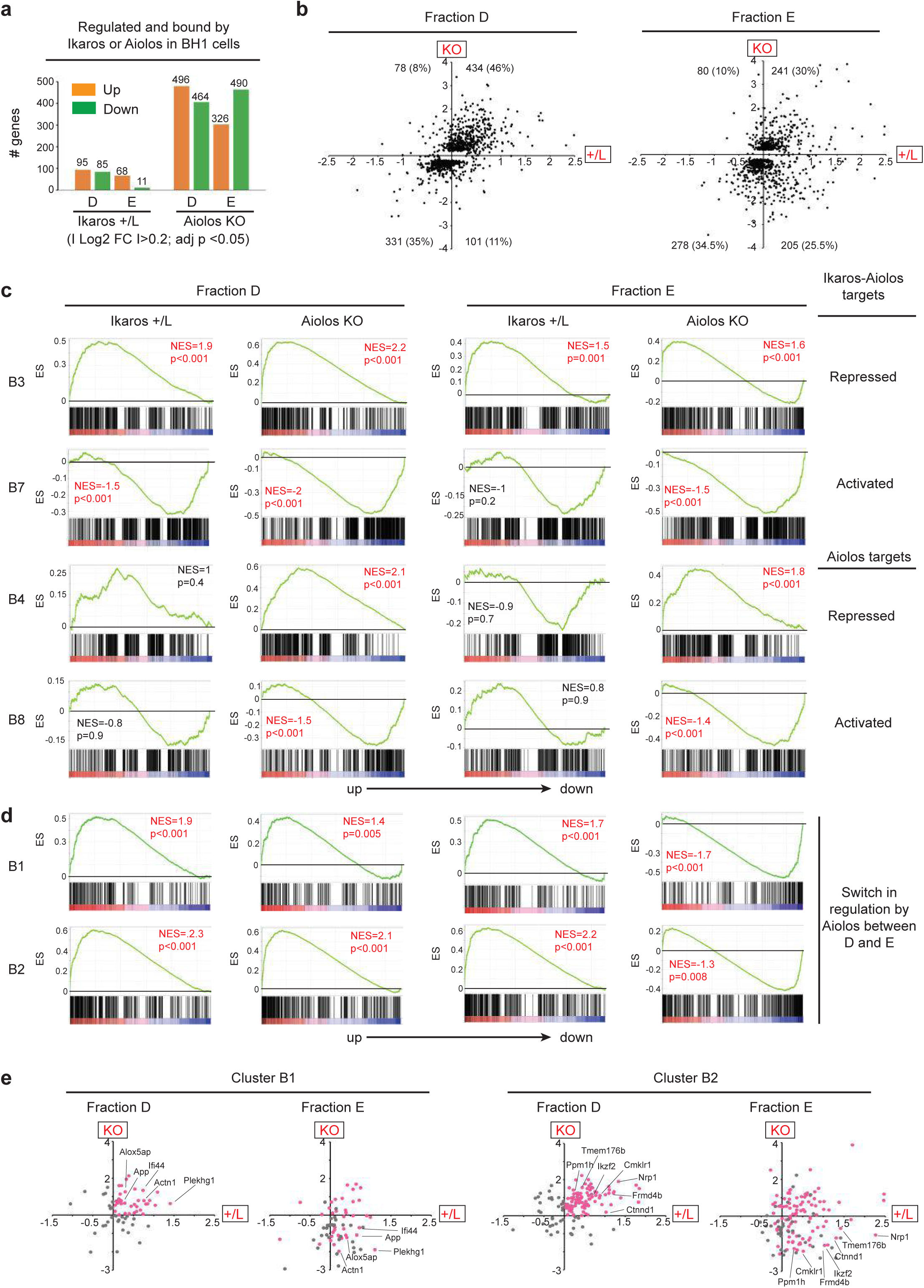
Requirement of Ikaros and Aiolos in small preB and immature IgM+ B cells. **(a)** Numbers of genes up- or downregulated in Ikaros +/L or Aiolos KO Fractions D and E cells. Selection criteria were: I Log2 FC I >0.2; adj p<0.05. **(b)** Scatter plots showing the Log2 FC of the genes deregulated in Fraction D or E, in Ikaros +/L (x-axis) and Aiolos KO (y-axis) cells. For each subtype, the genes represented are those deregulated in either Ikaros +/L or Aiolos KO cells. The number and proportion of genes in each quadrant are indicated. **(c, d)** GSEA plots showing enrichments of the genes from the indicated BH1 clusters among genes deregulated in Ikaros +/L or Aiolos KO Fraction D or E. Ranked genes comprise all genes detected in the RNA-seq analysis. ES: enrichment score. NES: normalized enrichment score. The p-value indicates the proportion of better enrichments obtained with 1000 random permutations of the ranked list. Red annotations highlight the GSEA plots showing significant enrichments. **(e)** Scatter plots showing the Log2 FC of cluster B1 or B2 genes that were detected as significantly changed in either Fraction D or E cells from Ikaros +/L or Aiolos KO samples (I Log2 FC >0.2 I; adj. p<0.05). Genes upregulated (Log2 FC>0) in both Ikaros +/L and Aiolos KO Fraction D cells are colored in pink. Names of selected genes with opposite responses in Fractions D and E from Aiolos KO mice are indicated.

To examine how individual target genes were affected by the Ikaros +/L and Aiolos KO mutations, we plotted the DETGs according to their response to TF LOF (Fig. 5b). In Fraction D, most of the DETGs were commonly up- or downregulated by the loss of either factor, consistent with a major role for the Ikaros-Aiolos heterodimers in small preB cells. In addition, the upregulated genes were more differentially expressed than the downregulated genes, indicating a strong repressive function of the heterodimers at this stage. However, the same was not true in Fraction E, where about half of the targets upregulated by Ikaros deficiency were instead downregulated by the Aiolos KO mutation. Thus Aiolos activates gene expression independently of Ikaros in Fraction E.

We then asked if the Aiolos-dependent DETGs from the BH1 system were also uniquely regulated by Aiolos in Fractions D and/or E, by using GSEA to assess the behavior of the genes from the various BH1 clusters in the mutant cell fractions (Figs. 5c and S10b). This revealed that the genes commonly repressed (cluster B3) and activated (B6, B7) by Ikaros and Aiolos (see Figs. 3d and S10b), were up- and downregulated, respectively, in the expected direction in the mutant subsets, particularly in Fraction D. In contrast, the genes specifically repressed (B4) and activated (B8) by Aiolos-Aiolos, were respectively enriched among the up- and downregulated genes only in the Aiolos KO fractions. Thus, the majority of the DETGs required the same dimer types for their expression in vitro and in vivo.

Interestingly, the genes strongly repressed by the Ikaros homodimers (clusters B1 and B2, corresponding to 22% of the DETGs), though faithfully enriched among the upregulated genes in the Ikaros +/L fractions, displayed strikingly different patterns of enrichment in the Aiolos KO fractions (Fig. 5, d and e). In Fraction D, B1 and B2 genes were upregulated in both of the mutant samples, indicating that Ikaros and Aiolos cooperate in repressing gene expression as this stage; given the low binding capacity of the Ikaros homodimers, this cooperation likely comes from the heterodimers. However, in Aiolos KO Fraction E cells, ∼50% of these targets were downregulated, even as they were still upregulated in Ikaros +/L cells. These results revealed a shift to Aiolos- Aiolos regulation in immature B cells, likely due to increasing Aiolos levels at this stage, and uncover a previously unrecognized antagonism between Aiolos-Aiolos activation and Ikaros- Ikaros repression.

Collectively, our results provide evidence that the homo- and heterodimers are differentially required to regulate hundreds of genes and multiple biological pathways at sequential steps of B cell differentiation. The Ikaros homodimer is uniquely present between early proB and large preB cells, and acts mainly as a repressor of genes that are shut off at early developmental stages. The Ikaros-Aiolos heterodimer is responsible for the largest number of gene expression changes in small preB cells, both as a repressor and activator. The Aiolos homodimer contributes to gene activation in immature IgM+ B cells, where it is required in part to counteract the repression exerted by the Ikaros homodimer.

### Aiolos rescues Ikaros null preB cell differentiation

Our data suggested that Aiolos is more potent than Ikaros in preB cells: Aiolos-containing dimers bind to a broader range of genomic targets than those containing only Ikaros, and Aiolos LOF disrupts the expression of a larger number of genes than Ikaros. We therefore hypothesized that Aiolos could rescue the block in preB cell differentiation caused by Ikaros LOF, and tested the ability of the dimers to rescue BH1 cell differentiation. The effects of low vs. high dox treatment on cells expressing Ikaros-Ikaros were first evaluated. High dox treatment led to the rapid downregulation of the λ5 surrogate light chain (SLC) at day 1 and the upregulation of the BCR by day 4 (Fig. S11, a and b), while low dox rescued preB cell differentiation with slower kinetics, as expected and in line with our previous results ^11^, indicating nonsaturating conditions. We then studied the effects of low dox treatment on all of the cell lines. The BH1 cells were dox-treated to express Ikaros-Ikaros, Ikaros-Aiolos or Aiolos-Aiolos, and analyzed over 5 days. Remarkably, Aiolos-Aiolos promoted a more rapid SLC downregulation and BCR upregulation after 1 and 2 days of treatment, respectively (Fig. 6, a and b), which was more potent than Ikaros-Aiolos, or Ikaros-Ikaros induced after low or high dox treatment. Aiolos-Aiolos expression was accompanied by a loss of cell viability (Fig. 6c, % DAPI-negative cells), due probably to the inability of the differentiated B cells to survive in these IL-7-supplemented conditions, while Ikaros-Aiolos expression resulted in an intermediate phenotype. These results demonstrated that ectopic Aiolos homodimers can rescue the differentiation of Ikaros KO preB cells better than Ikaros. Thus, in contrast to the highly specific roles of Ikaros and Aiolos at later steps of B cell differentiation, gene regulation normally driven by Ikaros at earlier stages can be functionally replaced by Aiolos.

**Figure 6.**
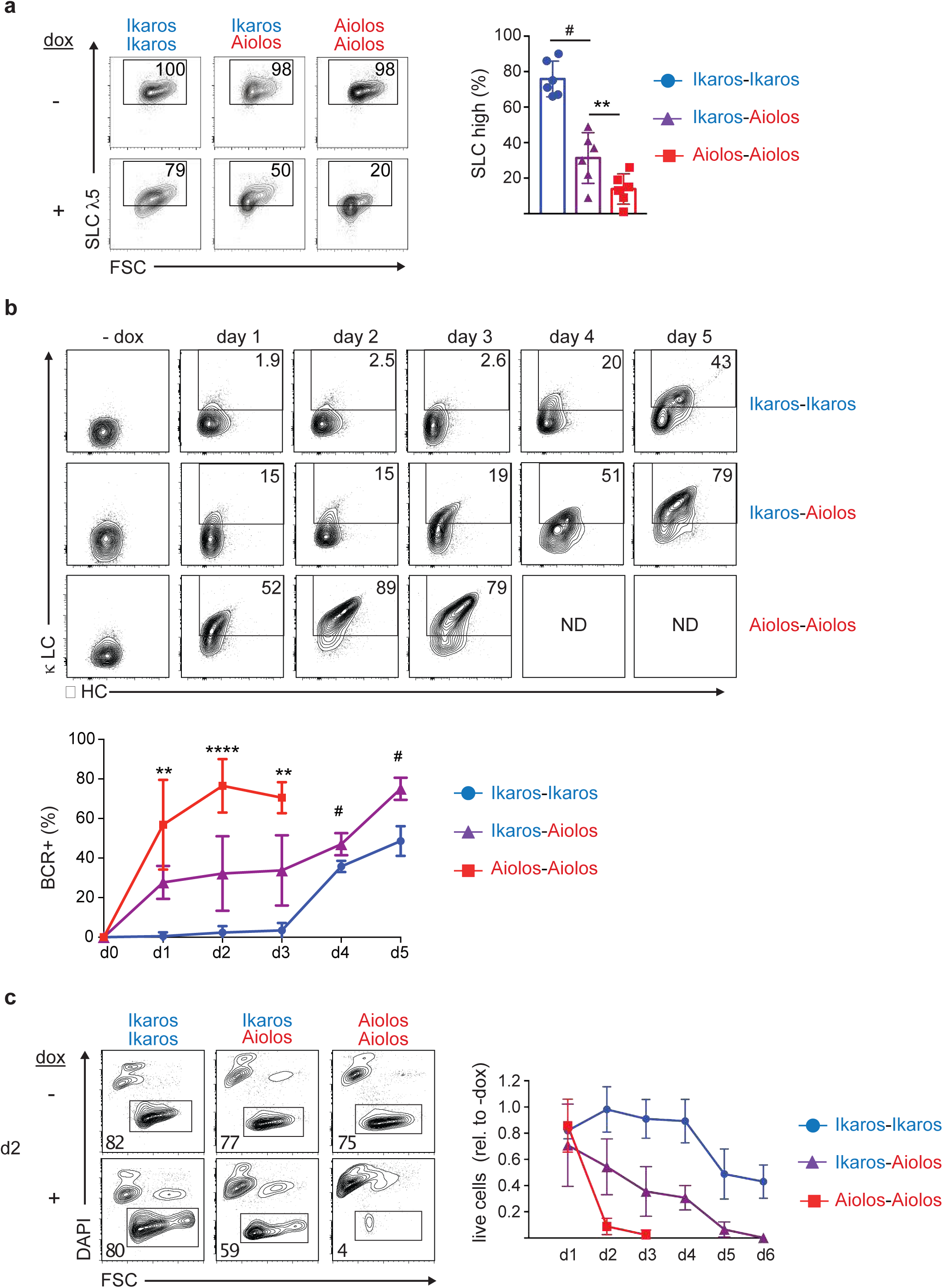
Aiolos-containing dimers rescue Ikaros null preB cell differentiation. In all panels, the indicated BH1-derived cells were induced to express the indicated proteins in low dox conditions. **(a)** Left: Representative plots for the expression of the surrogate light chain (SLC) λ5, after 24 h of dox. Right: Mean % of SLC hi cells after 24 h of dox (n=6 experiments). **(b)** Top: Representative plots for the expression of the BCR+ cells in culture, as monitored by the expression of the κ light (LC) and μ heavy chains (HC) (gated on live cells). For Aiolos-Aiolos cells, no data were collected for d4 and d5, as there were no surviving cells at these timepoints. Bottom: Mean % of BCR+ cells at days 1-5 (average of 6 experiments per sample). **(c)** Left: Cell viability after 2 d of dox treatment, as analyzed by DAPI staining and the forward scatter (FSC). Right: Mean % of live cells relative to cells not treated with dox, over 6 d of culture (average of 6 experiments per sample). ^#^p<0.05 comparison of Ikaros-Ikaros vs. Ikaros-Aiolos. **p<0.01; ****p<0.0001 (comparison of Aiolos-Aiolos vs. Ikaros-Aiolos) by Student’s t test.

### Ectopic Aiolos promotes the differentiation and cell cycle exit of murine BCR-ABL^+^ BCP-ALL cells

That ectopic Aiolos can complement Ikaros deficiency in preB cell differentiation suggested that it may do the same for leukemic cells. We first compared our RNA-seq data with lists of genes repressed or activated by Ikaros re-expression in mouse and human BCR-ABL+ BCP-ALL cells carrying *Ikzf1* LOF mutations ^41,42^. Indeed, there was a significant enrichment of BH1 genes repressed or activated by Ikaros and Aiolos (Fig. 7, a and b), including *Ctnnd1* and *Ifitm3*, whose high mRNA levels in BCP-ALL cells have been linked to disease severity.

**Figure 7.**
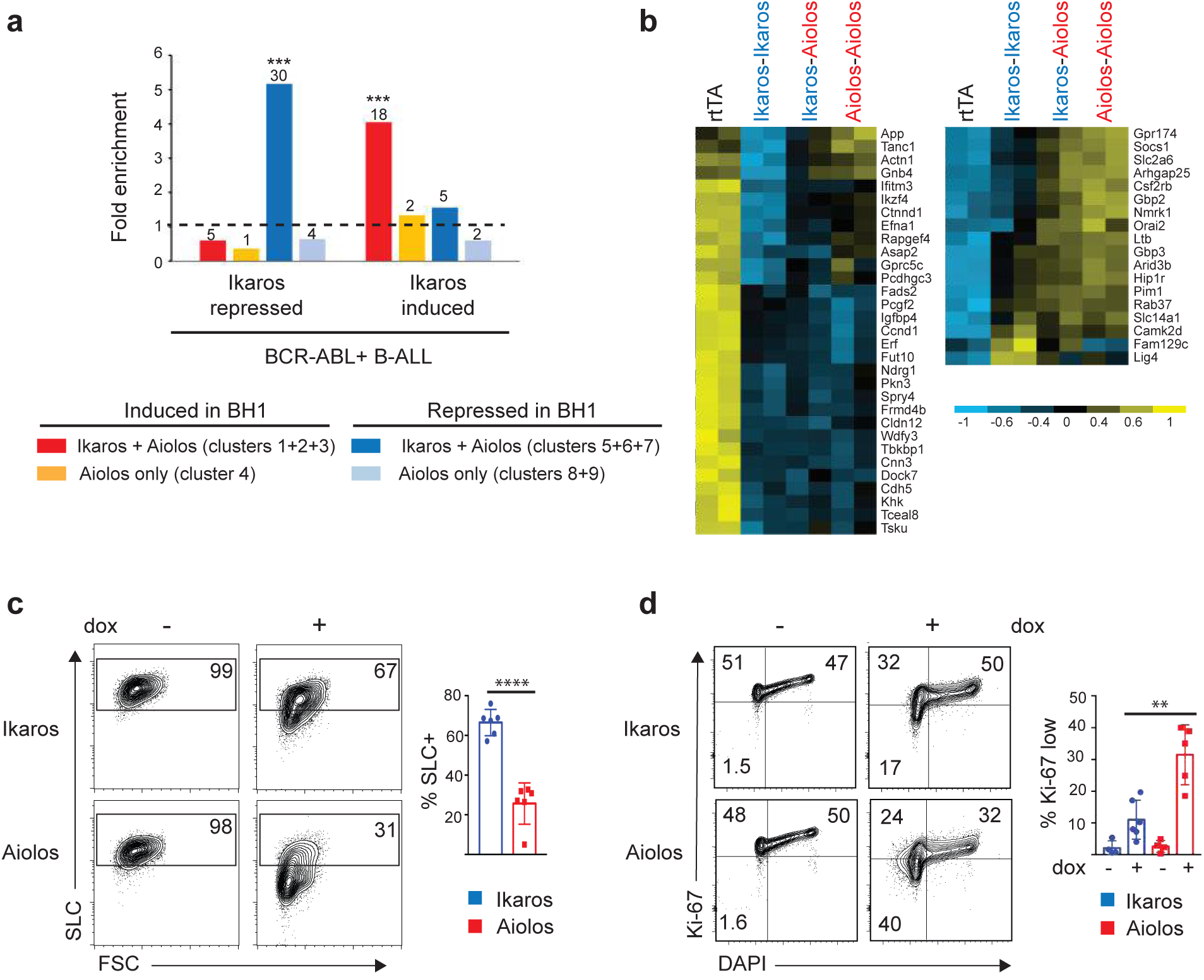
Ectopic Aiolos promotes the differentiation and cell cycle exit of murine BCR-ABL+ BCP-ALL cells. **(a)** Enrichment of genes repressed or activated by Ikaros and/or Aiolos in BH1 cells among the genes repressed or activated by Ikaros in the context of murine and human BCR-ABL1+ BCP-ALL cells (genes listed in Fig. 4 of Schjerven et al ^41^ and Fig. 3 of Witkowski et al ^50^). ***p<10^-8^ (hypergeometric test). **(b)** Heatmap of the expression of the genes repressed (left) or activated (right) by Ikaros in BCR-ABL1+ BCP-ALL and in BH1 cells. **(c, d)** BH1-BCR-ABL1 cells were transduced to express dox inducible Ikaros and Aiolos and GFP. GFP+ cells were analyzed after 48 h of high dox. (c) Left: Representative staining of intracellular SLC λ5 and forward scatter (FSC). Right: Mean % of SLC hi cells (n=6 experiments). (d) Left: Representative cell cycle analysis showing Ki-67 and DAPI levels as a measure of DNA quantity. Right: Mean % of Ki-67 low cells (n=5 experiments). **p<0.01; ****p<0.0001 (Student’s t-test).

To determine if Aiolos can counterbalance the effects of Ikaros LOF in Ikaros null BCR- ABL^+^ BCP-ALL cells, we studied BH1 cells transduced with the BCR-ABL oncogene. BCR-ABL^+^ BH1 cells were blastic and IL7-independent (not shown). The cells were further transduced to express dox-inducible Ikaros or Aiolos, along with a GFP reporter. High dox treatment for 2 days revealed that Aiolos promoted leukemic cell differentiation as measured by the downregulation of SLC λ5, and cell cycle exit as measured by a reduction in Ki-67 positivity, better than Ikaros (Fig. 7, c and d). However, Aiolos expression in the leukemic cells did not trigger the loss of cell viability that we observed in non-transformed BH1 cells. These results indicated that ectopic Aiolos has anti-tumor activity in Ikaros-deficient leukemic cells.

## DISCUSSION

The ability to dimerize is prevalent among transcription factor families, but how dimerization affects gene regulation was mostly unknown. Here we report the first direct study of Ikaros family homo- and heterodimers, and show that Ikaros and Aiolos are not equivalent. Ikaros-Ikaros, Ikaros-Aiolos and Aiolos-Aiolos bind distinct target sequences resulting in differential gene regulation. They are essential for different biological pathways, and required at sequential steps of B cell differentiation (see graphical abstract).

One of the most striking discoveries was the difference in the size of the target gene repertoire associated with each dimer. Ikaros homodimers are directly required for the regulation of relatively few genes, a particularly unexpected result given that Ikaros is a prominent differentiation factor and tumor suppressor. Indeed, selective Ikaros-Ikaros targets are never detected, and all Ikaros-Ikaros target genes are recognized by the Aiolos-containing dimers. Ikaros homodimer binding is strongest at genes important for early B cell differentiation, like those responsible for IL7 signaling, and cell adhesion and migration, where we and others have shown that Ikaros is important for repressing the expression of genes activated by IL7/STAT5 signaling and for promoting cell migration away from IL7-rich zones ^11,14,36^. In contrast, Ikaros-Aiolos and Aiolos-Aiolos bind vastly more target genes which are linked to processes important for late B cell differentiation, B cell receptor signaling, and various immune response pathways such as the NFkB pathway. The ability of the Aiolos-containing dimers to promote BCR and NFkB signaling may explain why *IKZF3* GOF mutations are frequently detected in chronic lymphocytic leukemias, which are associated with a high expression of BCR signaling genes and strong NFkB activity ^31,32^. Ikaros-Aiolos and Aiolos-Aiolos also exhibit distinct binding patterns, where the heterodimers bind with lower, or higher, intensities to specific targets. This is especially true at targets where Ikaros homodimer binding is markedly different from that of the Aiolos homodimers (see *Lrrc74a* and *Phkg2* in Fig. 1e), and heterodimer binding is skewed toward the dominant partner. Thus, Aiolos defines the size of the target gene repertoire, but binding strength is modulated by both subunits.

The difference in binding is due largely to the ability of Aiolos to recognize more nucleotide combinations immediately flanking the core GGAA motif. Another difference is that the Aiolos- biased motifs preferentially lack a T immediately 3’ of the GGAA, which is a feature of the commonly bound motifs. Q183, in ZF3 of the DNA binding domain, is required for characteristic Aiolos binding. This position is occupied by an arginine in Ikaros and all other family members, as well as in Ikaros homologs of non-vertebrate deuterostomes; R183 is also present in ancestral Aiolos proteins but replaced with a glutamine in tetrapods. Why Aiolos Q183 was selected for in tetrapods is intriguing. Tetrapods comprise the only evolutionary branch that make the conventional B2 cells studied here, which are essential for adaptive antibody responses and immunological memory ^43^. The R183 to Q change in Aiolos might have evolved at the same time as the B cell lineage, perhaps to help establish the B2 cell gene expression program. Recent studies have reported that patients with immune dysregulation ^44,45^, or therapy-resistant multiple myeloma ^46^, show *IKZF1* GOF mutations that substitute the canonical R183 for cysteine or histidine, associated with stronger DNA binding. Conversely, our experiments demonstrated that mutating Q183 to arginine in Aiolos results in a loss of binding to Aiolos-biased sites. Thus, we propose that Q183 is required for the more permissive binding of Aiolos to DNA.

A second unexpected finding was that the dimers are functionally different. Ikaros homodimers act mainly as transcriptional repressors while Aiolos homodimers function broadly as activators, and the Ikaros-Aiolos heterodimers as either repressors or activators. Whether a common target is repressed or activated may be determined by local dimer concentration. For example, *Igll1* and *Myc* are bound by all dimers, but repressed most efficiently by Aiolos-Aiolos. Numerous other genes are coregulated by Ikaros and Aiolos, including similarly repressed genes that were originally identified as Ikaros targets in other cell types (eg. *Ctnnd1*, *Hes1*, *Tmem176b*, *Rhoj*, *Dock1*, *Cd9*) ^47–49^, as well as genes important for the tumor suppressive function of Ikaros ^41,50^. The latter group may correspond to a core program that is repressed by Ikaros or Aiolos regardless of dimer or cell type.

Our results illustrate the important and sequential roles of Ikaros and Aiolos in B cell development. The ubiquitous presence of Ikaros, together with the increase of Aiolos in small preB and immature B cells, imply a shift in equilibrium from Ikaros-Ikaros in proB and large preB cells (Fractions B to C/C’), to Ikaros-Aiolos in small preB cells (Fraction D), and Aiolos-Aiolos in immature IgM+ B cells (Fraction E). Indeed, the genes repressed by the Ikaros homodimers are downregulated in Fractions B and C/C’ cells, while the cooperation between Ikaros and Aiolos, likely via heterodimerization, plays a considerable role in establishing the gene expression program in Fraction D cells. This cooperation is reduced in Fraction E cells, where Aiolos-Aiolos antagonizes Ikaros-Ikaros repression at a subset of target genes involved in cell adhesion and locomotion. Thus, both Ikaros and Aiolos are essential for B cell differentiation in the bone marrow, and their orchestrated assembly into functionally distinct dimer pairs imparts a critical layer of specificity to gene regulation, both in target gene selection and in activating vs. repressive activity.

This study sheds light on how Ikaros and Aiolos mutations may contribute to the loss of dimer function in human pathologies. Ikaros, but not Aiolos, LOF are frequent in BCP-ALLs with a proB cell phenotype, in line with its exclusive expression at this stage, and the essential function of the Ikaros homodimers in regulating early B cell differentiation and tumor suppressor gene networks. Notably, the Aiolos homodimers work better than the Ikaros homodimers to induce the cell cycle exit of murine BCP-ALL cells, a phenotype that is probably linked to the ability of Aiolos to potently repress *Myc*. These results suggest that Aiolos expression might be a promising therapeutic avenue for BCP-ALL treatment, especially for leukemias with Ikaros haploinsufficiency. Future investigation will be required to assess the potential effect of Aiolos as an anti-cancer agent.

Loss of homo- or heterodimer function may also underlie Ikaros family dysfunctions in B cell immunodeficiencies. Ikaros LOF mutations are linked to early B cell developmental defects and loss of mature B cell numbers with age; they likely affect Ikaros homodimer activities, and possibly those of the Ikaros-Aiolos heterodimers, at various B cell stages ^25^. Similarly, an Aiolos G159R mutation leads to early B cell developmental defects ^24^; these proteins appear to act in a dominant-negative manner by forming dysfunctional heterodimers with Ikaros. Another Aiolos mutation (N160S) just one amino acid away is associated with normal to elevated numbers of B cells that exhibit extensive functional abnormalities ^26^; it does not appear to interfere with Ikaros function, and may therefore affect Aiolos homodimer binding in mature B cells. Our new knowledge of gene regulation by Ikaros and Aiolos homo- and heterodimers will help dissect how these different mutations contribute to disease.

In conclusion, our study has revealed how combinations of Ikaros family dimers regulate widespread gene expression changes during B cell development. It will be important to determine if Ikaros and Aiolos act similarly in other cell types, and how dimerization with other family members influences binding specificity. The split-epitope approach will be an invaluable tool to dissect the function of this and other transcription factor families, whose activities are regulated by dimerization choice.

## MATERIALS AND METHODS

### Recombinant proteins

The mouse Ikaros and Aiolos sequences used in this study correspond to NM_001025597 (Ikaros) and NM_011771.1 (Aiolos). The V1 and V2 fragments of the Venus protein (respectively amino acids 2-158 and 159-239) were linked C-terminally to the Ikaros and Aiolos proteins with a serine/glycine-rich linker (SGSGGGGSGGGGSSG). IkΔDD proteins were deleted for the C-terminal amino acid segments 454-482 and 490-504.

### Cell lines

The non-transformed, IL7-dependent BH1 *Ikzf1* null preB cell line was described previously ^11^. BH1 cells and their derivatives were cultured in IMDM supplemented with 10% FCS (tetracycline-free for cell lines expressing the rtTA protein, Dutcher), 10% supernatant from the IL7-producing cell line J558-IL7 ^51^, 1% penicillin/streptomycin and 50 μg/ml gentamycin.

The BH1 lines expressing the inducible V1 and V2 fragments were generated after 3 consecutive retroviral transductions. First, BH1-rtTA cells expressing the rtTA transactivator were produced (pMSCV-rtTA-PGK-Neo vector, also carrying the neomycin resistance gene), and neomycin selected (1 mg/ml). Second, inducible constructs for the Ikaros- or Aiolos-V2-tagged proteins were inserted [TtlCPH-TRE3G-PGK-puro vector ^52^], to carry the sequences under the control of the tetracycline response element TRE3G, and the cDNA for the puromycin N-acetyltransferase under the control of the PGK promoter, and puromycin selected (3 μg/ml). Third, the V1-tagged constructs were integrated (TtlCPH-TRE3G-PGK-blasticidin), and blasticidin selected (20 μg/ml). Doxycycline-dependent expression of the V1- and V2-tagged constructs was verified by flow cytometry, Western blot and fluorescence microscopy. Note that the cell lines used to map Aiolos dimer binding with Mab199 by ChIP-seq expressed an Aiolos(Δ2-23)-V2 protein instead of full-length Aiolos. The N-terminal deletion did not affect DNA binding, as no differences in Aiolos binding were found when compared with an independently generated cell line expressing full-length Aiolos-V1 and -V2 proteins (anti-Aiolos ChIP-seq, not shown).

BH1 cells expressing BCR-ABL1 were generated by retroviral transduction of BH1 rtTA cells with pMSCV-BCR-ABL1-IRES-CD8 (also expressing the truncated human CD8 as marker), and sorted for CD8+ cells. These cells were then transduced with the TtlCPH-Ikaros-IRES-GFP- PGK-puro or TtlCPH-Aiolos-IRES-GFP-PGK-puro vector and puromycin selected.

### Mice

The L hypomorphic *Ikzf1* allele was described previously ^38^. The *Ikzf3* null allele was generated by CRISPR/Cas9-mediated deletion of the sequences between the chr11:98466968 and chr11:98467878 genomic positions (mm10), via electroporation of appropriate gRNAs and the Cas9 protein into fertilized eggs (C57BL/6N). This deletion comprises the 3’ part of intron 7 and the coding sequence of exon 8. Mice were bred in an SPF facility and studied at 6-8 weeks of age according to IGBMC Ethical Committee (Com’Eth) guidelines.

### Electrophoretic mobility shift assays

Protein extracts (3 μg of nuclear extracts from Cos cells transfected with the appropriate expression vector) were first preincubated for 30 min on ice with 0.5 μg poly dIdC (Merck-Sigma) in a total volume of 20 μl of binding buffer (20 mM Hepes pH 7.9, 0.2 mM EDTA, 20% glycerol, 100 mM KCl, 1 mM DTT). ^32^P-labeled probes (100 fM in 2 μl) were then added, incubated on ice for another 30 min before electrophoresis on a 5% polyacrylamide gel and autoradiography. For antibody supershifts, 0.5 μl of the Mab199 Ab (0.75 ng) was added to the binding reaction 5 min before loading on the gel. The BS4 and *Cish* probes were described previously ^7,36^. Probes derived from Aiolos specific targets were (core GGAA motifs underlined):

*Tnfrsf13c*: 5’ GCCAGGCAGGAAGTGAAAACGGCCTGCAGGAAGCACC;

*Sh2b3*: 5’ TCCAGGACTGGCTCACAAGGAAGCCAGGGG;

*Ighj4*: 5’ TGGCAGGAAGCAGGTCATGTGGCAAGGCTATTTGGGGAAGGGA.

Probes derived from the Sh2b3 or Ighj4 backbones were described in the figures and text. Quantification of the signals from bound complexes was done with the ImageJ software.

### RNA sequencing

WT B cell BM subsets were sorted from 6- to 8-week-old C57BL/6 mice as B220+CD19+CD43+IgM-BP1- (Fraction B), B220+ CD19+ CD43+ IgM- BP1+ (Fraction C/C’), B220+CD19+CD43-IgM- (Fraction D), and B220+CD19+CD43-IgM+ (Fraction E) cells. Aiolos KO and Ikaros +/L BM subsets were sorted as B220+CD43-IgM- (Fraction D) and B220+CD43-IgM+ (Fraction E) cells. RNA was extracted from 1-4x10^6^ cells, using the RNeasy Plus Mini (Qiagen) or Nucleospin RNA (Macherey-Nagel) kits. Aiolos KO or WT littermates were 6-10 weeks old, and Ikaros +/L mice and WT littermates 7 weeks old. RNA from BH1-derived cells were prepared from 10^6^ cells. Libraries were prepared with the TruSeq Stranded mRNA kit (Illumina), and sequenced (HiSeq 4000; Illumina) with single-end 50 bp read length. Reads were pre-processed to trim the first base in samples prepared with the Stranded mRNA Prep Ligation protocol to remove adapters (Cutadapt v1.10), poly(A) and low-quality sequences (Phred quality score <20) in all samples. Reads <40 bases were discarded, and then aligned to the M. musculus genome mm10 (STAR v2.5.3a). Gene expression quantification was performed from uniquely aligned reads (HTSeq v0.6.1p1), with annotations from ENSEMBL release 90 and “union” mode. Read counts were normalized with the median-of-ratios method ^53^. Differential gene expression analyses were performed using the Bioconductor package DESeq2 (v1.16.1) on R (v3.3.2). P-values were adjusted for multiple testing ^54^. Clustering analysis was performed with Gene Cluster 3.0 and heatmaps generated by Java TreeView. Pathway analyses were performed with Metascape (http://metascape.org). GSEA was performed using GSEA 2.0 ^55^.

### ChIP and ChIP-sequencing

For ChIP-seq experiments with splenic B cells, cells were isolated [biotinylated anti-CD19 antibody, followed by streptavidin-coated magnetic beads (Miltenyi Biotec)]. Protein crosslinking from 50x10^6^ cells were performed in 2 mM disuccinimidyl glutarate (ChemCruz) for 45 min at room temperature (RT). Protein-DNA crosslinking was performed in 1% paraformaldehyde (PFA) for 10 min at RT followed by addition of 2 M glycine for 5 min at RT to quench. Cell lysis was performed in the presence of a protease inhibitor cocktail (Roche #11836153001) as follows: 10 min at 4°C with LB1 buffer (50 mM tris-HCl pH8, 2 mM EDTA, 0.1% NP-40, 10% glycerol) followed by 10 min at 4°C with pre-chilled LB2 buffer (10 mM tris-HCl pH8, 1 mM EDTA, 0.5 mM EGTA, 200 mM NaCl). Nuclei were resuspended in LB2.3 (10 mM tris-HCl pH8, 1 mM EDTA, 0.5 mM EGTA, 200 mM NaCl, 0.033% Na-deoxycholate, 0.16% N-laurylsarcosine) and sonicated for 6 cycles (5 min high, 15 sec on, 15 sec off) with a Bioruptor 200 (Diagenode), to yield 150-300 bp fragments. The sonicated chromatin was diluted 10x with a dilution buffer (50 mM tris-HCl pH8, 200 mM NaCl, 5 mM EDTA, 0.5% NP-40) and pre-cleared for 1 h at 4°C with appropriate sepharose beads (80 μl of 50% bead slurry for 1 ml of diluted chromatin). For anti-Ikaros or anti-Aiolos ChIP, the cleared chromatin was incubated ON at 4°C with rabbit polyclonal anti-Ikaros (in-house) or rabbit monoclonal anti-Aiolos (Cell Signaling #15103). IP was performed with protein A-Sepharose beads (Sigma) in the presence of 0.1% SDS for 1 h at 4°C. V1-V2 complexes were IP-ed with the mouse monoclonal Ab Mab199 (SS and DvE, to be published elsewhere), which was previously crosslinked to protein G sepharose beads by dimethyl pimelimidate (20 μg of Ab for 60 μl of 50% bead slurry, 30 min at RT; Abcam protocol for crosslinking Abs to beads). IP-ed complexes were washed 6x in washing buffer (20 mM tris-HCl pH8, 500 mM NaCl, 2 mM EDTA, 1% NP-40, 0.1% SDS) and 3x in TE, then eluted in 2% SDS in TE (10 mM tris-HCl pH8, 1 mM EDTA) for 1 min at RT. De-crosslinking was performed in the elution buffer ON at 65°C. Input chromatin was de-crosslinked in 200 mM NaCl in the same conditions. DNA was purified with phenol/chloroform extraction. Libraries were prepared using the MicroPlex Library Preparation kit v2 (Diagenode #C05010014). Single-end 50 bp sequences were obtained with the HiSeq 4000. Image analysis and base calling were performed using RTA 2.7.7 and bcl2fastq 2.17.1.14. Adapter-dimer reads were removed using DimerRemover (https://sourceforge.net/projects/dimerremover/). The reads were aligned to mm10 using Bowtie 2 v2.3.4.3. Reads were filtered with respect to the quality of the alignment to mm10 using Samtools View v1.9 (MAPQ=40). Read numbers were the following: Ikaros-Ikaros low dox (LD): 30.4x10^6^, high dox (HD): 37.2x10^6^; Aiolos-Aiolos LD: 39.8x10^6^, HD: 43.1x10^6^; Ikaros-Aiolos LD: 31.5x10^6^, HD: 30.5x10^6^. All analyses were performed with a subset of 15x10^6^ reads per sample. Peak identification was performed with MACS2 Callpeak v2.1.1.20160309.6. Peak annotation was done with Homer. Heatmaps were generated with SeqMINER ^56^. Differential analyses were done with DEseq2 v1.16.1, considering the low and high dox samples for a given dimer as replicates. To generate the boxplot in Fig. 1c, we first extracted the genomic positions of regions having peaks in all datasets using Bedtools Intersect v2.30.0 ^57^. Then, the positions of peak summits for Aiolos_homodimer_HD were extracted (Bedtools Intersect). Total numbers of mapped reads were scaled down to 15x10^6^ for all samples and read size was extended to the expected fragment length of 200. Finally, the number of fragments per sample at summit positions was calculated (Bedtools Intersect). Plots were created using in-house R scripts with R v4.1.1 and ggplot2 v3.3.5. For Fig. S4e, the numbers of fragments at the union of merged peaks per condition (Ikaros-Ikaros, Ikaros-Aiolos, Aiolos-Aiolos) were calculated (Bedtools Intersect), and the plots created using R v4.1.1 and ggplot2 v3.3.5.

### Motif analysis

The 4 nt sequences corresponding to the -2, -1, +1 and +2 nt positions before and after the GGAA sequences within a 90 bp window under the peak summits were extracted manually and the motifs associated with these sequences generated with *ggseqlogo* ^58^.

### Flow cytometry and cell sorting

The following reagents were used: anti-B220-PE-Cy7, anti-CD19-PerCP-Cy5.5, anti-CD43- BV421 or -PE, anti-CD24-biotin, anti-κ light chain-biotin, anti-λ light chain-biotin, anti-CD179- biotin or -BV421, anti-BP-1-BV421 (all BD Biosciences); anti-B220-FITC and SA-Alexa Fluor405 (Invitrogen); anti-Aiolos-PE or -eF660 (eBioscience); anti-Ki67-AlexaFluor 700 (Biolegend); anti-IgM-Cy5 (Southern Biotech), and anti-rabbit-PE (Jackson ImmunoResearch). For cytoplasmic protein staining, the Fix and Perm kit (eBioscience) was used; for nuclear protein staining, the Foxp3/transcription factor Fix/Perm kit (eBioscience) was used. Dead cells were excluded by DAPI or Zombie AquaV430 (Biolegend). Cells were analyzed with the FACS Celesta or FACS LSR II analyzers (BD Biosciences) and the FlowJo software (TreeStar, Ashland, OR). Cell sorting was done with FACS Aria II or FACS Aria Fusion cell sorters (BD Biosciences); purity was >95%.

### Microscopy

7x10^4^ cells were deposited on a Superfrost Plus slides (Thermofisher), then fixed with 4% PFA, and permeabilized with 0.5% triton-X100. Ikaros staining with the anti-Ikaros Ab was revealed with a goat anti-rabbit secondary antibody coupled to the AF568 (Fisher scientific SAS).

### Immunoprecipitation

HEK 293T cells were transfected with the indicated plasmids using Lipofectamine 2000. After 24 h, whole cell extracts were prepared by lysing the cells in 500 μl of lysis buffer (20 mM tris-HCl pH 7.5, 150 mM NaCl, 5 mM EDTA pH 8.0, 5 mM NaPiP, 1 mM Na3VO4, 20 mM NaPO4 pH 7.6, 3 mM b-glycerophosphate, 10 mM NaF,1% Triton X-100, supplemented with complete protease inhibitor cocktail (Roche #11873580001) and phosphatase inhibitor cocktail 3 (Sigma #P0044) for 10 min on ice. Lysates were sonicated using the Bioruptor 200 (15 sec on, 15 sec off, high power) for 15 min at 4°C. Supernatants were pre-cleared at 13K rpm for 10 min. 30 μg was kept for the input. IP was performed on 2 mg of proteins using 8 μg of Mab199) and protein G sepharose (30 μl) ON. Protein complexes were washed 3x in lysis buffer, separated by SDS-PAGE and subjected to Western blotting using the anti-Ikaros antibody.

## ACKNOWLEDGEMENTS

We thank Johannes Zuber (Institute for Molecular Pathology, Vienna) for the tetracycline-inducible retroviral vectors, members of the Chan-Kastner lab for scientific discussions and help. We thank the IGBMC GenomEast platform for help with sequencing, Céline Keime for help with RNA-seq data analysis, the IGBMC flow cytometry facility (C. Ebel, M. Philipps), the Institut Clinique de la Souris (Marie-Christine Birling) for mouse generation, the IGBMC animal facility (M. Gendron, S. Falcone, W. Magnant, A. Vincent), and the IGBMC cell culture facility.

S.C. and P.K. received funding from the Agence Nationale de la Recherche (ANR-17-CE15-0023-01, ANR-22-CE15-0017), the Fondation pour la Recherche Médicale (Equipe FRM 2019, EQU201903007812), an equipment grant from LNCC Grand Est/Bourgogne Franche Comté, institute funds from INSERM, CNRS, Université de Strasbourg. This work is part of the Interdisciplinary Thematic Institute (ITI) IMCBio program from the University of Strasbourg, CNRS and INSERM, which is supported by IdEx Unistra [(ANR-10-IDEX-0002), SFRI-STRAT’US (ANR-20-SFRI-0012) and EUR IMCBio (ANR-17-EURE-0023)]. M-C.D. was funded by predoctoral fellowships from the ANR-10-LABX-0030-INRT grant and the FRM, and postdoctorally by the ANR grant ANR-17-CE15-0023-01. A.A. was funded by predoctoral fellowships from IMCBio and Fondation ARC. Q.H. was funded by a post-doctoral fellowship from Fondation ARC. The GenomEast platform is a member of the France Génomique consortium (ANR-10-INBS-0009).

## Author contributions

Performed experiments: M-C.D., B.H., A.A., Q.H., P.M., C.C., P.K.

Analyzed data: M-C.D., B.H., A.A., S.L.G., I.M.L.B., D.v.E., P.K., S.C.

Provided the BiFC system, the Mab199 antibody and valuable advice: D.v.E., S.S. Supervised the work: B.H., P.K., S.C.

Designed the study: M-C.D., B.H., P.K., S.C.

Wrote the manuscript: P.K., S.C., and all authors reviewed

## Accession Number

ChIP-seq and RNA-seq data are available under the GEO accession number GSE2223.

