## Supplementary information for "How Ikaros and Aiolos homo- and heterodimers drive gene expression to ensure B cell development"

Supplementary Figure Legends
Suppementary figures S1 to S11

### SUPPLEMENTARY FIGURE LEGENDS

**Figure S1. Expression of Ikaros and Aiolos during B cell differentiation.**

Top: Gating strategy to identify BM B cell Fractions A-F. Middle: Flow cytometry analysis of Ikaros and Aiolos in the indicated fractions and splenic B cells. For Ikaros, control corresponds to the signal obtained with the secondary Ab alone (PE-coupled anti-rabbit IgG). For Aiolos, control corresponds to staining of Aiolos KO cells. Bottom: Relative mean fluorescence intensities (MFI) from 4 (Ikaros) and 8 (Aiolos) independent experiments. Data are relative to the MFI for Fraction F. *$p<0.05$; ***$p<0.001$; ****$p<0.0001$ (paired t-test).

**Figure S2. Labeling of homo or heterodimers with Venus.**

**(a)** Tagging of specific dimers with the Venus reporter. Each protein partner was tagged at the C-terminus with the V1 or V2 fragments. Dimerization leads to assembly of V1 and V2, reconstitution of Venus fluorescence, and generation of the epitope for the Mab199 antibody, which does not recognize the individual V1 or V2 domains.

**(b)** Representation of Ikaros and Aiolos tagged with V1 or V2, with the SGSGGGGSGGGGSSG linker to provide flexibility.

**(c)** Flow cytometry analysis of Venus expression in the indicated BH1 cell lines treated or not with high dox (24 h).

**(d)** Visualization of Venus by microscopy in the indicated cells treated with high dox over time. Nuclei were stained with DAPI.

**Figure S3. Tagging of Ikaros with V1 and V2 does not affect its function.**

**(a)** Tagged Ikaros dimers accumulate in discrete nuclear foci. BH1 cells expressing inducible untagged Ikaros or tagged Ikaros-Ikaros were cultured for 24 h with high dox, and analyzed by microscopy for Venus and Ikaros expression. Nuclei were stained with DAPI.

**(b)** Differentiation of cells induced to express Ikaros-Ikaros after 5 d in high dox, as measured by the surface expression of the light ($\kappa/\lambda$; LC) and heavy ($\mu$; HC) chains of the BCR.

**(c)** Analysis of Ikaros binding to its target site at the *Cish* gene (Cish 184; Heizmann et al., 2020), by ChIP with anti-Ikaros or Mab199, from cells induced to express Ikaros-Ikaros (high dox, 16 h). Cish 4200 corresponds to a region not bound by Ikaros located at +4200 bp of the *Cish* TSS.

**(d)** V1 and V2 by themselves do not induce dimerization. Full-length or dimerization domain (DD)-deleted Ikaros were tagged with V1 or V2, and expressed separately or together in 293T cells, as indicated. Mab199 was used for the IP.

**(e)** Tagged full-length complexes can be supershifted but not dimerization-deficient ones. Nuclear extracts from Cos cells transfected with the indicated constructs were tested by EMSA on the BS4 target site. Complexes were supershifted with Mab199.

**(f)** V1 and V2 do not increase the stability of DNA-bound Ikaros complexes. Tagged Ikaros-Ikaros or untagged Ikaros were bound to the BS4 probe, and stability of binding was investigated by adding a 100x excess of unlabeled probe for the indicated times. Open arrowheads indicate dimeric complexes, while closed arrowheads indicate higher order complexes.

**Figure S4. Dimer expression levels in low vs. high dox conditions.**

**(a)** Flow cytometry analysis of Venus expression in BH1 cells treated for 24 h with low or high dox. CTRL corresponds to cells expressing only V1-tagged proteins.

**(b)** Comparison of Ikaros and Aiolos levels in cells expressing the indicated tagged Ikaros and/or Aiolos vs. the physiological counterparts in BM immature IgM+ B cells (for Ikaros) or splenic CD19+ B cells (for Aiolos). CTRL corresponds to secondary Ab staining alone.

**Figure S5. Mapping DNA binding by specific dimers by ChIP-seq.**

**(a)** Genome browser tracks showing Ikaros and Aiolos homo- and heterodimer binding (Mab199 ChIP-seq) in BH1 cells expressing the indicated tagged dimers (high and low dox samples; top), as well as Ikaros and Aiolos binding in cells expressing Ikaros-V2 or Aiolos-V2 (high dox, anti-Ikaros and anti-Aiolos ChIP-seq; bottom) in a representative genomic region (Chr. 2: 27,100,000-27,500,000, mm10). Arrowheads point to peaks predominantly bound by Aiolos-containing dimers (open) or by all dimers (closed).

**(b)** Heatmaps of the peak intensities detected in the ChIP-seq experiments from cells expressing the indicated tagged Ikaros or Aiolos, using the indicated Abs. The Mab199 ChIP-seq data are from the high dox samples. $19x10^6$ reads for each sample were used to generate the heatmaps in (b) and (c).

**(c)** Heatmaps of the intensities of the peaks selected as being detected in both Ikaros-Ikaros and Aiolos-Aiolos ChIP-seq experiments (245 peaks from Fig. 1d, top panel).

These peaks were also similar in intensity when detected with anti-Ikaros or anti-Aiolos antibodies.

**(d)** Representative peaks that were bound similarly by Ikaros-Ikaros or Aiolos-Aiolos (left), and those bound specifically by Aiolos-Aiolos (right), when analyzed with Mab199, anti-Ikaros or anti-Aiolos (in cells expressing Ikaros-V2 or Aiolos-V2).

**(e)** Scatter plots of the peak intensities of low and high dox samples for the indicated dimer.

**(f)** Examples of target sites selectively bound by Ikaros-Aiolos and Aiolos-Aiolos (high dox), using Mab199. Aiolos binding in splenic CD19+ B cells is shown at the bottom, using anti-Aiolos.

**(g)** Heatmaps of binding of the indicated dimers in BH1 cells and Aiolos binding in splenic CD19+ B cells, using Mab199 (BH1) and anti-Aiolos (spleen).

**Figure S6. Differential DNA binding by Ikaros and Aiolos in vitro.**

**(a)** Sequence conservation of the indicated Aiolos-specific targets among mammals. GGAA motifs are boxed. Sequences used as EMSA probes are underlined.

**(b)** Dimeric binding of Aiolos to the *Sh2b3* probe. Nuclear extracts from Cos cells expressing the indicated proteins were used in the EMSAs. The complex bound by the Aiolos-Aiolos extract was supershifted by Mab199.

**(c)** Influence of flanking nucleotides on Ikaros and Aiolos binding. Probes were derived from the *Sh2b3* probe backbone. The two 5' and 3' nts adjacent to the core GGAA motif (green) were altered to match those found in the BS4 or *Cish* target sequences (red).

**(d)** Selection of motifs associated with common or Aiolos-biased peaks. 84 commonly bound and 132 Aiolos-biased peaks were selected manually. The nucleotides surrounding all GGAA (or TTCC) motifs present within 90 nts of the peak summits were collected and used for the analyses in (e) and Figs. 2, b and d.

**(e)** Distribution of the 4 nts in commonly bound or Aiolos-biased peaks at the -2, -1 and +2 positions. The p-values associated with the distribution differences were calculated with the Chi-square test, using the distributions seen with the common Ikaros and Aiolos peaks as references.

**(f)** Binding by the heterodimer to the indicated probes. Binding was performed with nuclear extracts from Cos cells expressing the indicated V1 and V2-tagged proteins, and binding by the dimer of the V1 and V2-tagged proteins was detected by supershift with Mab199 (arrowhead). Left: Representative EMSA experiment. Right: Quantification of 3 independent experiments. *p<0.05 (Student's t-test).

**Figure S7. Structural properties of Ikaros and Aiolos DBDs and their target sequences.**

**(a)** Plot of the roll angles over the 8 nt sequences for Aiolos-biased or common target sequences, as shown in the inset. The common sequences show large positive roll angles for the 5' flanking region of the GGAA core sequence (red ellipse), in line with the larger bending propensity of the YR 5' flanking sequence. In contrast, the Aiolos-biased sequences have small negative roll angles including the 5' flanking region (blue ellipse). Roll angles were calculated with the Deep DNAshape webserver [1].

**(b)** Positions of Ikaros R183 and Aiolos Q183 in an Alphafold3 model of the Ikaros and Aiolos DBD bound to DNA.

1. Li, J., and Rohs, R. (2024). Deep DNAshape webserver: prediction and real-time visualization of DNA shape considering extended k-mers. Nucleic Acids Res *52*, W7-W12. 10.1093/nar/gkae433.

**Figure S8. Regulation of Ikaros and Aiolos-bound genes in BH1 cells.**

**(a)** Representative genome browser tracks for genes from the gene sets used in Fig. 3d. For peak intensity comparisons, all vertical scales were set at 100. Note that *Cst7* and *Pkn3* were repressed by Ikaros or Aiolos, while *Nod2*, *Scn11a*, *Nabp1* and *Socs3* were activated by Aiolos.

**(b)** Genome browser RNA-seq tracks showing representative genes with distinct regulatory patterns.

**(c)** Proportions of genes from the B1 to B8 clusters strongly bound by Ikaros-Ikaros and Aiolos-Aiolos [left; peaks labeled as (++; ++) in (a) and Fig. 3c] or strongly bound by Aiolos-Aiolos but not Ikaros-Ikaros [right; peaks labeled as (+++; -) in (a) and Fig. 3c] (see legend of Fig. 3c for selection criteria). The dotted red line represents the proportion of bound genes expected in random gene pools selected from all genes detected in the RNA-seq experiment. Statistics were calculated with the Chi-square test, comparing the expected and observed gene numbers in each highlighted group. This analysis showed significant enrichment for common Ikaros-Ikaros and Aiolos-Aiolos binding among Ikaros-repressed genes (clusters B1-B3), but not among genes specifically repressed by Aiolos (B4), or activated by any dimer (B5-B8). It also shows significant enrichment for strong Aiolos-biased binding among genes displaying Aiolos-dependent activation (clusters B6-B8), but not among those specifically activated by Ikaros (B5), or repressed by any dimer.

**(d)** Proportions of genes bound by Ikaros-Ikaros, or only by Ikaros-Aiolos or Aiolos-Aiolos, among genes from the B1-B8 BH1 clusters. The green lines correspond to the expected proportion of bound genes if binding were equally distributed among all regulated genes. Statistics were calculated with the Chi-square test, comparing the expected and observed gene numbers in each highlighted group. This analysis showed a significant and specific enrichment for Ikaros-Ikaros binding among genes specifically activated by Ikaros (cluster B5). Aiolos-biased binding was also higher among genes displaying Aiolos-specific repression (B4), though only marginal significance was reached.

**Figure S9. Ikaros and Aiolos-dependent genes during B cell differentiation.**

**(a)** Enrichment numbers for the genes from the BH1 cell clusters B1-B8 that were found among the genes up- or downregulated during the developmental transitions between the indicated fractions (schematized in Fig. 4a). The total gene pool that was considered comprises 15,145 genes that were detected in the RNA-seq experiments for both the BH1 and the primary cell populations (genes annotated as "Gene models" or pseudogenes were excluded). The expected gene number for the intersection between cluster y and developmental transition z corresponds to $(n_y/15{,}145) \times (n_z)$. Enrichments correspond to the ratios between the observed and expected gene numbers. P-values were calculated with the hypergeometric test. Dark-shaded areas highlight cluster/developmental transition pairs for which enrichments were >2, while light-shaded areas highlight pairs with lower but significant enrichments (1.5-2).

**(b)** Pathway enrichment among the 102 genes downregulated by Ikaros in BH1 cells (clusters B1 or B2) and also downregulated between Fractions B and C/C' (Metascape). The 31 genes belonging to enriched pathways related to cell adhesion or cell locomotion are represented.

**Figure S10. Analysis of Aiolos KO and Ikaros +/L mice.**

**(a)** BM B cell populations in Aiolos KO mice. Left: Representative flow cytometry analysis of the indicated B cell fractions. Right: Mean % of the indicated fractions in

WT vs. KO BM (n=5 experiments, 6- to 8-week-old mice). *p<0.05; ***p<0.001 (Student's t-test).

**(b)** GSEA plots showing enrichments of the genes from the B5 and B6 BH1 clusters among genes deregulated in Ikaros +/L or Aiolos KO Fraction D or E cells. Ranked genes comprise all genes detected in the RNA-seq analysis. ES: enrichment score. NES: normalized enrichment score. The p-value indicates the proportion of better enrichments obtained with 1000 random permutations of the ranked list.

**Figure S11. Effects of low and high dox on the differentiation of BH1 cells re-expressing Ikaros.**

**(a)** Left: Representative levels of SLC $\lambda$5 among Ikaros-Ikaros BH1 cells cultured with no, low or high dox for 24 or 48 h. Right: Mean % of SLC hi cells (n=6 experiments for low dox; n=3 experiments for high dox).

**(b)** Left: Representative proportion of BCR+ cells among Ikaros-Ikaros BH1 cells cultured with no, low or high dox for 4 or 5 d. Right: Mean % of BCR+ cells (n=4 experiments for low dox; n=2 experiments for high dox). **p<0.01; ***p<0.001 (Student's t test).

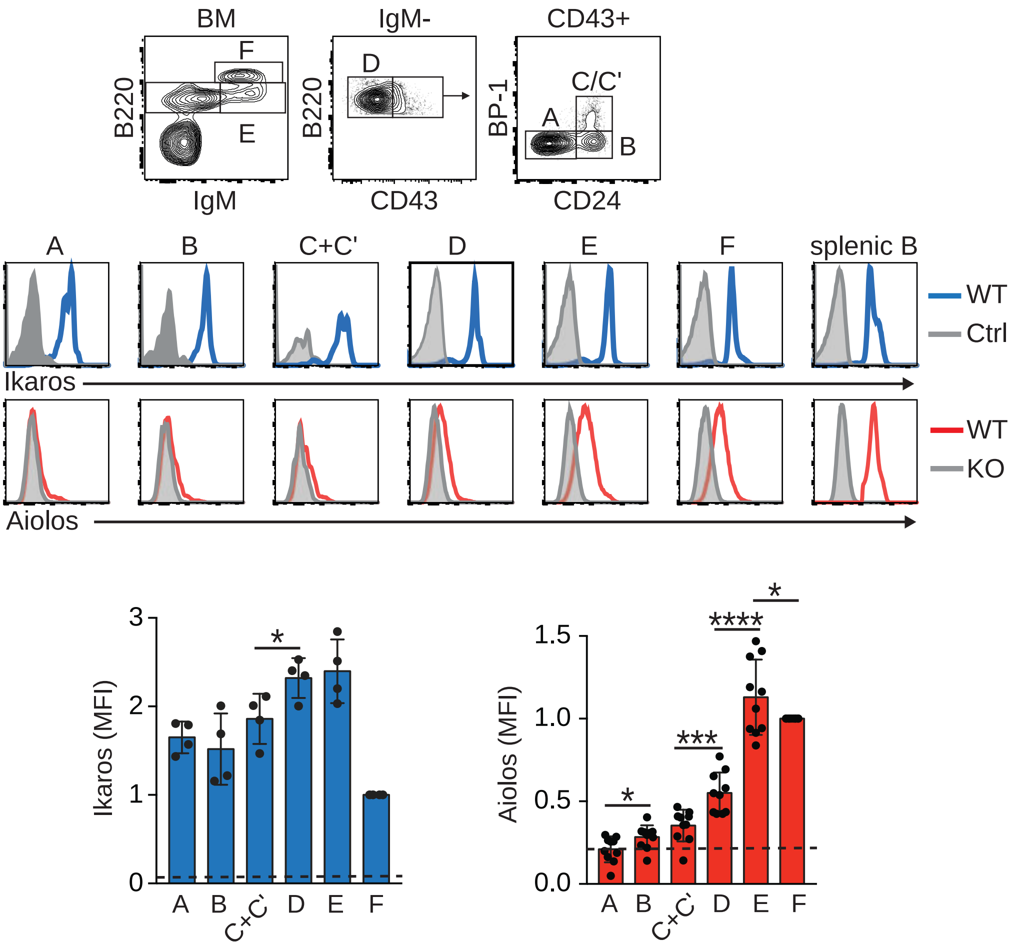


Figure S1
Deau et al

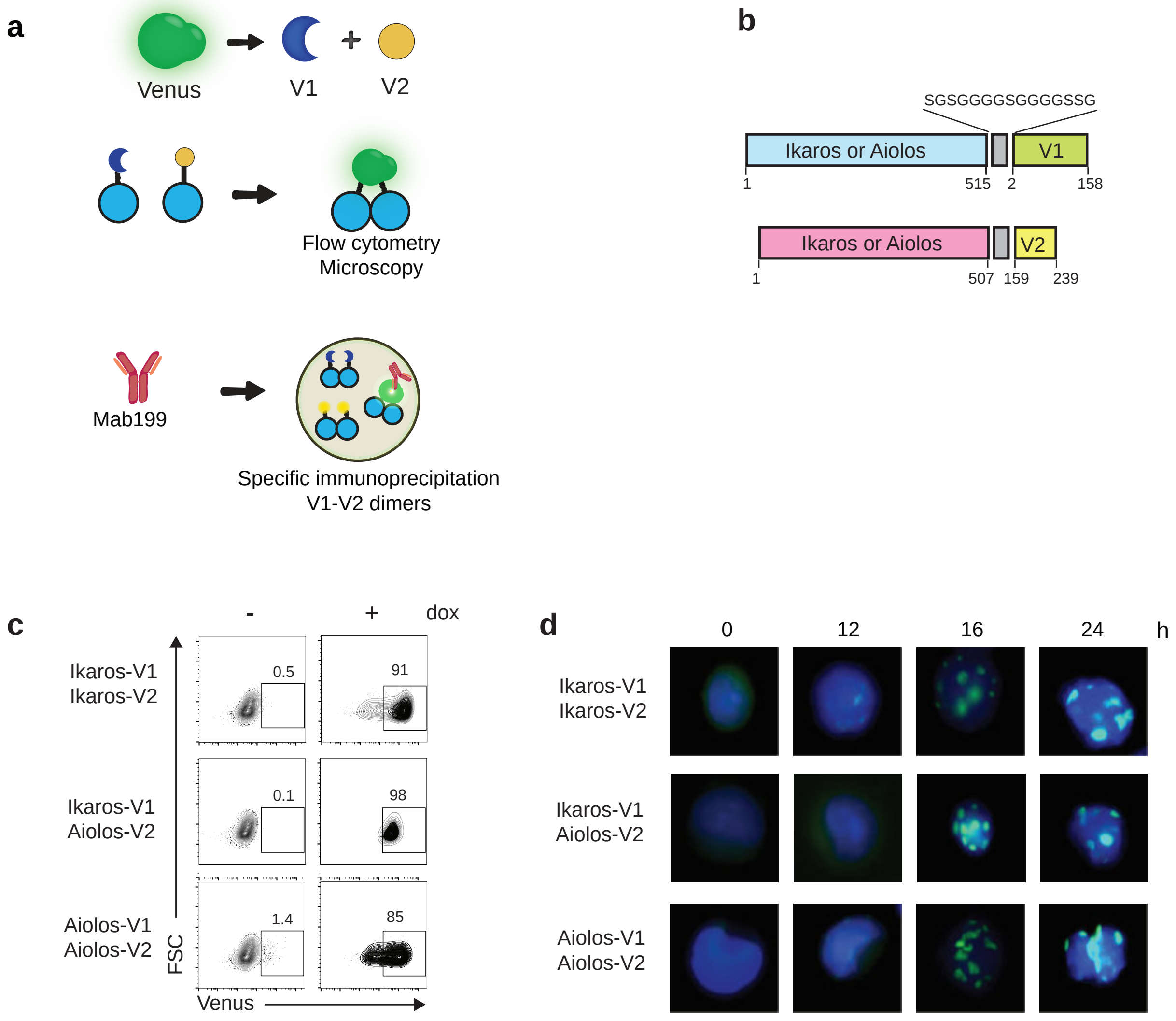


Figure S2
Deau et al

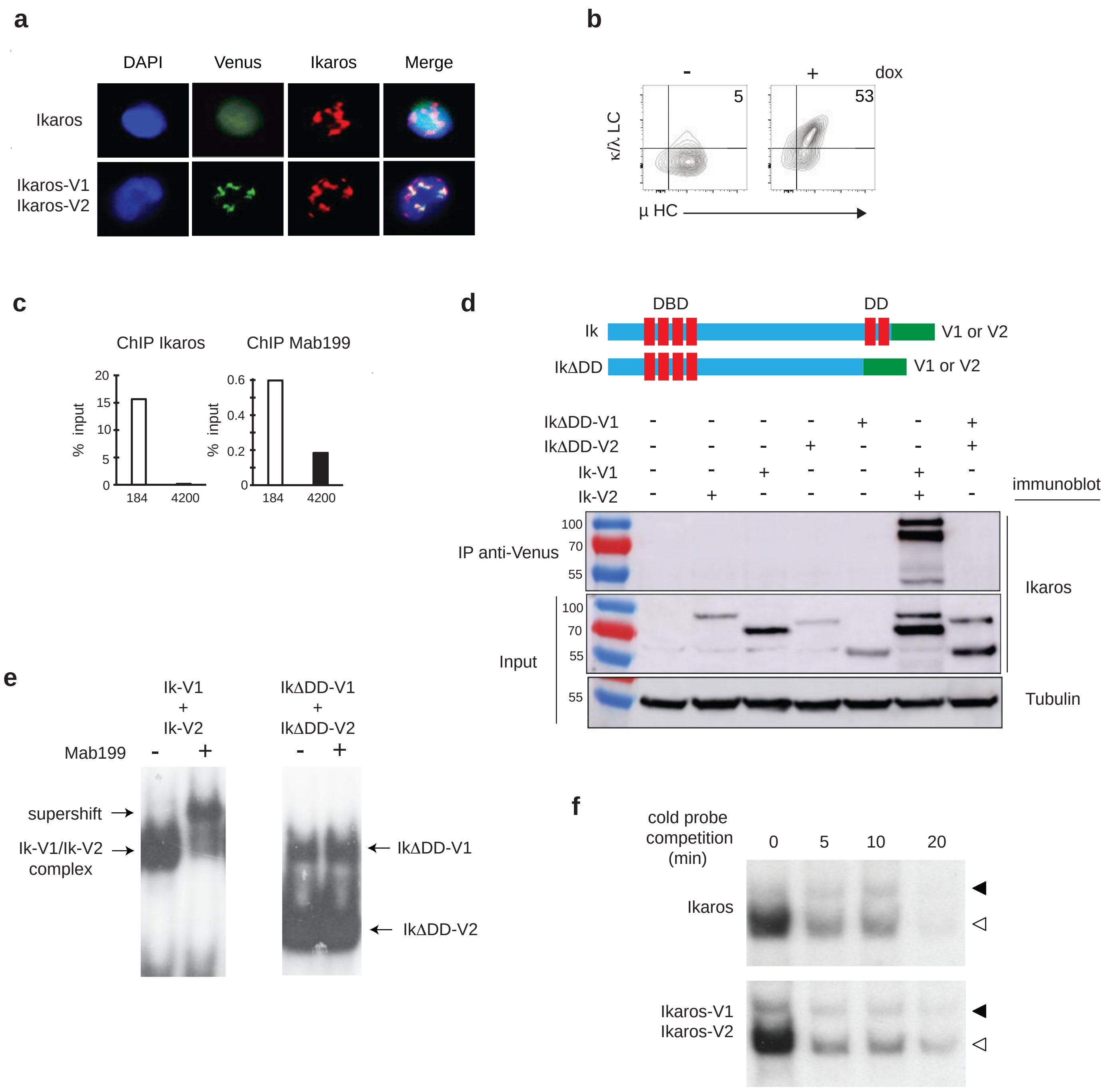


Figure S3
Deau et al

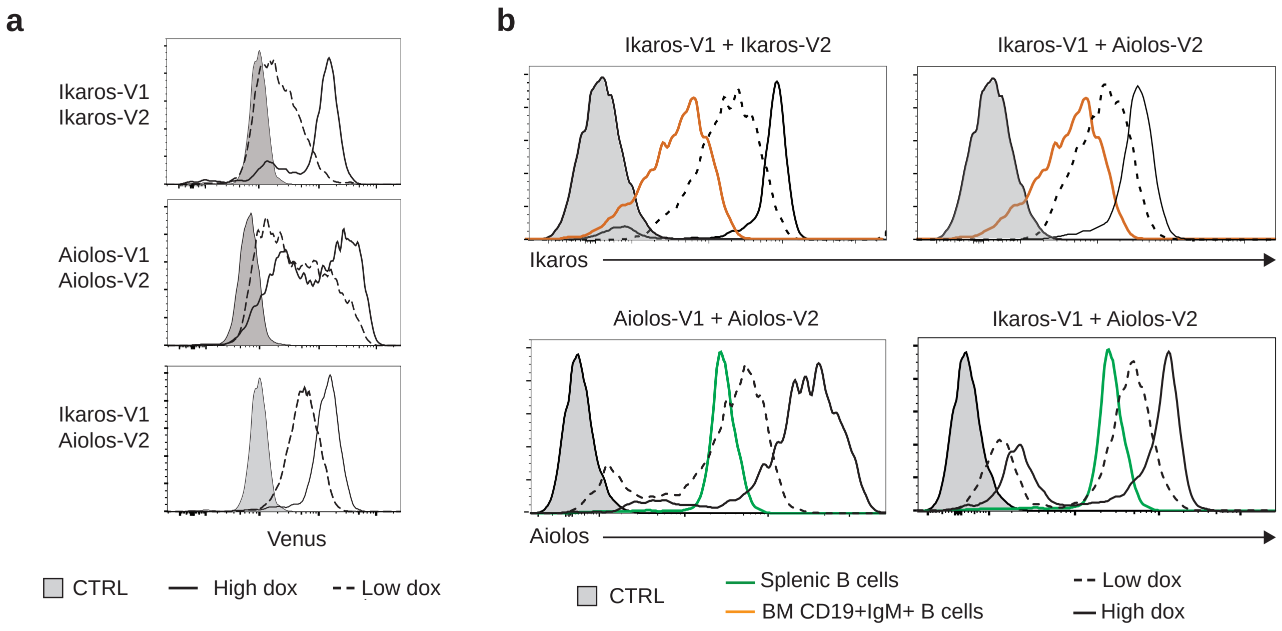


Figure S4
Deau et al

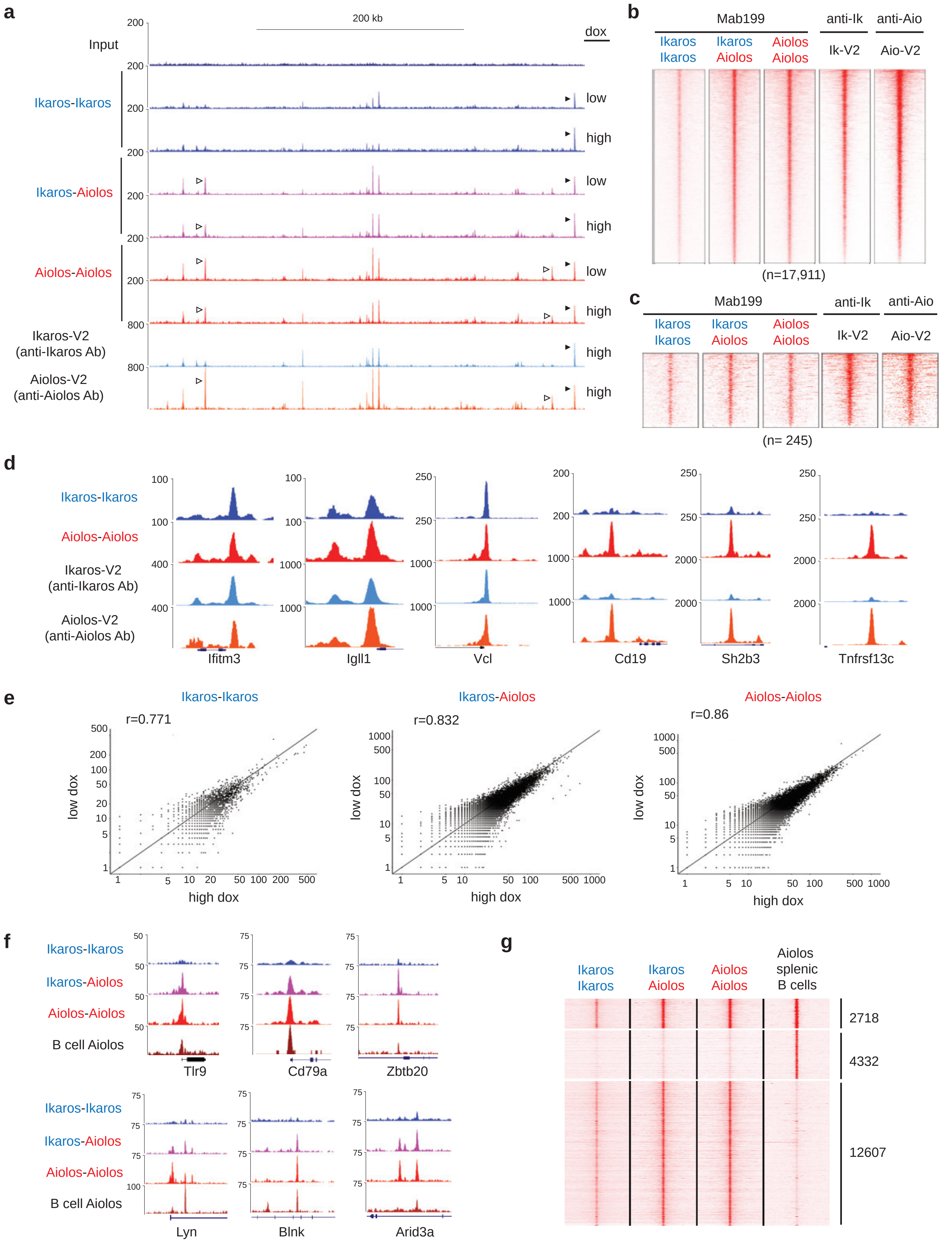


Figure S5
Deau et al

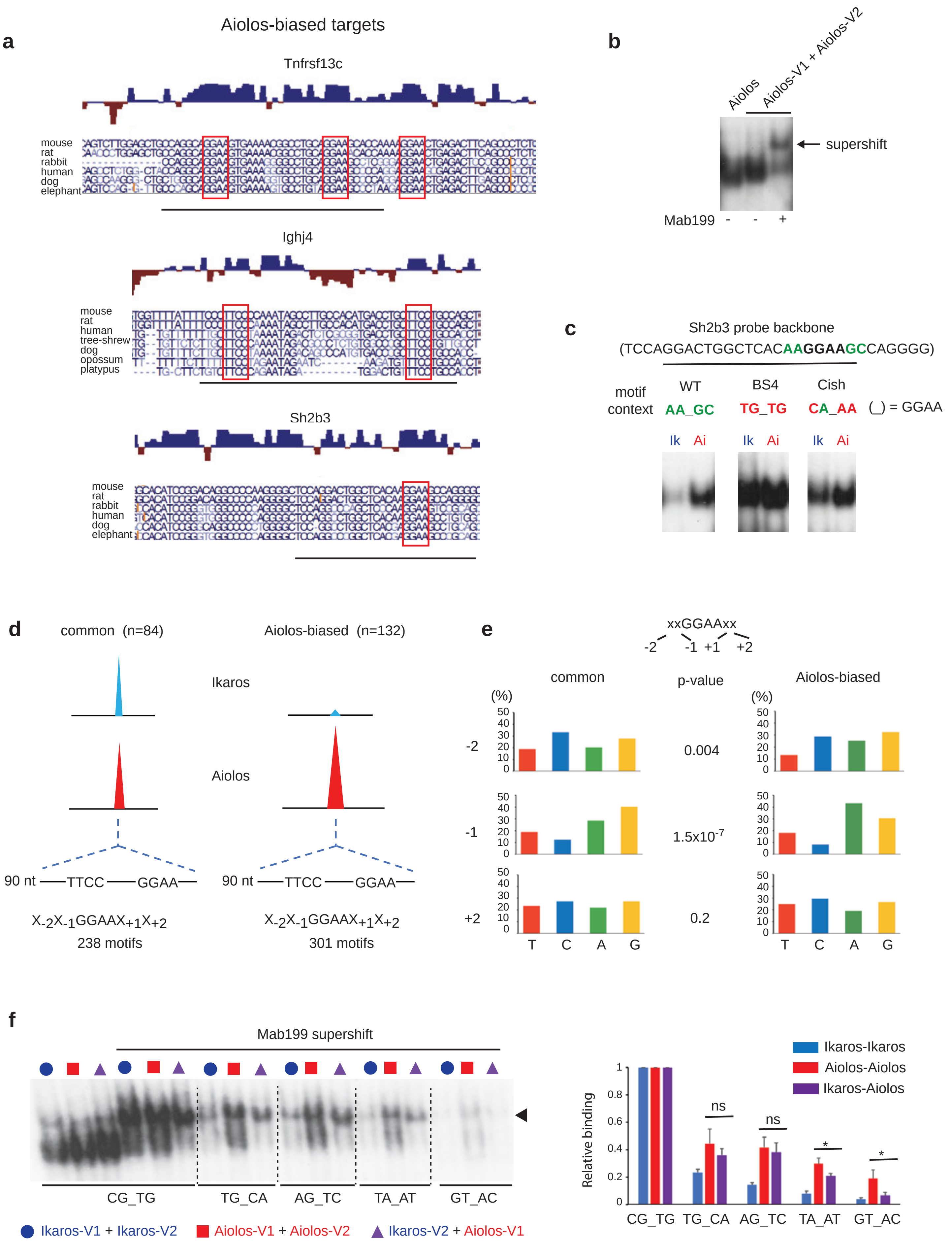


Figure S6
Deau et al

**a**

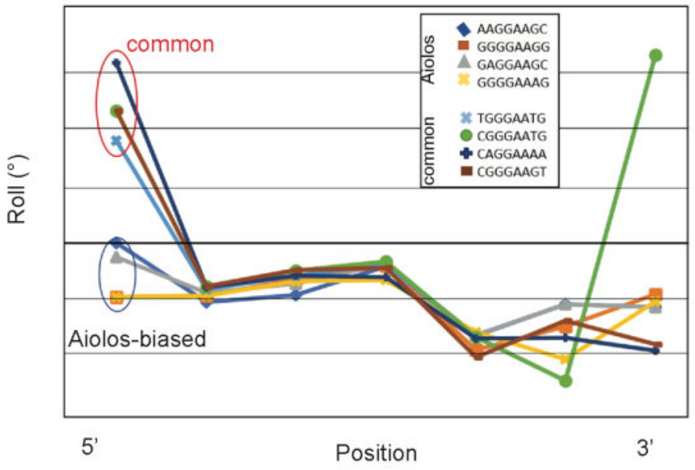


**b**

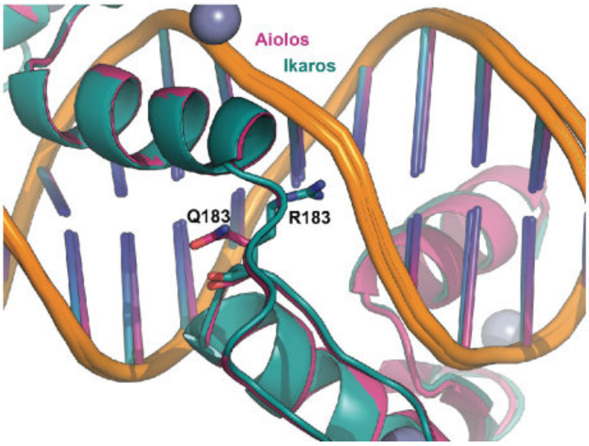


Figure S7
Deau et al

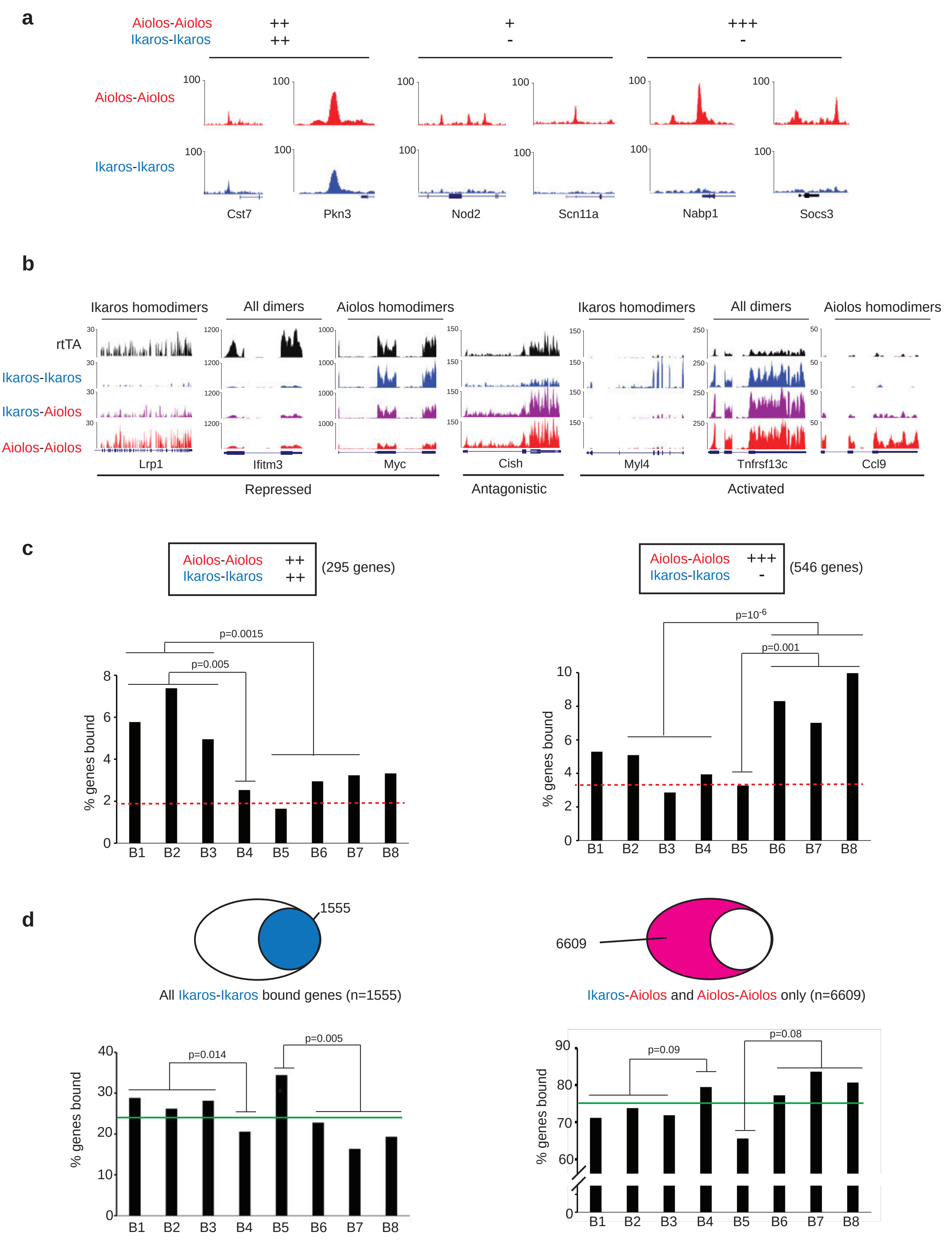


Figure S8
Deau et al

**a**

| # genes observed | # genes expected | enrichment | p-value |
|---|---|---|---|

Downregulated genes

| | B to C/C' (840) | | | | C/C' to D (2020) | | | | D to E (1959) | | | |
|---|---|---|---|---|---|---|---|---|---|---|---|---|
| B1 (203) | **42** | **11** | **3.73** | **$6x10^{-14}$** | 33 | 27 | 1.21 | 0.03 | 40 | 26 | 1.52 | $6.8x10^{-4}$ |
| B2 (383) | **60** | **21** | **2.8** | **$1.5x10^{-13}$** | **104** | **51** | **2.03** | **$1.7x10^{-13}$** | 84 | 49 | 1.69 | $2.7x10^{-7}$ |
| B3 (381) | 24 | 21 | 1.12 | 0.07 | **136** | **51** | **2.6** | **$3.7x10^{-29}$** | **116** | **49** | **2.34** | **$9.7x10^{-20}$** |
| B4 (355) | 4 | 19 | 0.2 | $10^{-5}$ | **168** | **47** | **3.5** | **$6.2x10^{-56}$** | 65 | 46 | 1.41 | $8x10^{-3}$ |

Upregulated genes

| | B to C/C' (496) | | | | C/C' to D (1593) | | | | D to E (1824) | | | |
|---|---|---|---|---|---|---|---|---|---|---|---|---|
| B5 (61) | **10** | **2** | **4.9** | **$2x10^{-5}$** | **20** | **6.5** | **3.06** | **$2.2x10^{-6}$** | 9 | 7.5 | 1.2 | 0.12 |
| B6 (369) | 12 | 12 | 1 | 0.15 | **131** | **39** | **3.37** | **$8.4x10^{-39}$** | **113** | **44** | **2.54** | **$3.9x10^{-22}$** |
| B7 (552) | 6 | 15 | 0.33 | $5x10^{-4}$ | **226** | **58** | **3.89** | **$10^{-81}$** | **113** | **66** | **1.7** | **$3x10^{-9}$** |
| B8 (329) | 7 | 10 | 0.64 | 0.07 | **116** | **34** | **3.35** | **$4.2x10^{-34}$** | **95** | **39** | **2.39** | **$6x10^{-17}$** |

**b**

| | | B1 | B2 | Cell substrate adhesion | Actin filament based process | E cadherin stabilization paythway | Positive regulation of cell motility | Rho signaling pathway | Cdc42 GTPase cycle |
|---|---|---|---|---|---|---|---|---|---|
| Abca1 | ATP binding cassette subfamily member 1 | ■ | | | | | | ■ | |
| Abi2 | Abl interactor 2 | ■ | | | ■ | | | | ■ |
| Dock9 | Dedicator of cytokinesis 9 | ■ | | | | | | ■ | |
| Gab1 | Grb2 associated binding protein 1 | ■ | | | | | ■ | | |
| Glipr2 | GLI pathogenesis related 2 | ■ | | | | | ■ | | |
| Iqsec2 | IQ motif and Sec7 domain ArfGEF 2 | ■ | | | ■ | | | ■ | |
| Map3k12 | Mitogen-activated protein kinase kinase kinase 12 | ■ | | | | | ■ | | |
| Plin3 | Perilipin 3 | ■ | | | | | | | ■ |
| Sema7a | Semaphorin 7A | ■ | | ■ | | ■ | | | |
| Specc1 | Sperm antigen with calponin homology and coiled coil domains 1 | ■ | | | ■ | | | | |
| Zyx | Zyxin | ■ | | ■ | ■ | ■ | | | |
| Abr | ABR activator of RhoGEF and GTPase | | ■ | | ■ | | | | ■ |
| Arhgap5 | Rho GTPase activating protein 5 | | ■ | ■ | | ■ | ■ | | ■ |
| Cd40 | CD40 molecule | | ■ | ■ | | | ■ | | |
| Coro2b | Coronin 2B | | ■ | | | ■ | | | |
| Dock6 | Dedicator of cytokinesis 6 | | ■ | | | | | ■ | ■ |
| Emilin1 | Elastin microfibril interfacer 1 | | ■ | ■ | | ■ | ■ | | |
| Fhl3 | Four and a half LIM domains 3 | | ■ | | ■ | | | | |
| Frmd6 | FERM domain containing 6 | | ■ | | ■ | | | | |
| Igf1r | Insulin like growth factor 1 receptor | | ■ | ■ | | ■ | ■ | | |
| Inf2 | Inverted formin 2 | | ■ | | ■ | | | | |
| Itga5 | Integrin subunit alpha 5 | | ■ | ■ | ■ | ■ | ■ | | |
| Lamb2 | Laminin subunit beta 2 | | ■ | ■ | | ■ | | | |
| Lpp | LIM domain containing preferred translocation partner in lipoma | | ■ | | | ■ | | | |
| Myh10 | Myosin heavy chain 10 | | ■ | | ■ | | | | |
| Ntn1 | Netrin 1 | | ■ | | | | ■ | | |
| Plxd1 | Plexin D1 | | ■ | | | | | | ■ |
| Sash1 | SAM and SH3 domain containing 1 | | ■ | | | | ■ | | |
| Stard8 | StAR related lipid transfer domain containing 8 | | ■ | | ■ | | | ■ | ■ |
| Tbxa2r | Thromboxane A2 receptor | | ■ | | | | | | ■ |
| Trio | Trio Rho guanine nucleotide exchange factor | | ■ | | | | | | ■ |

Figure S9
Deau et al

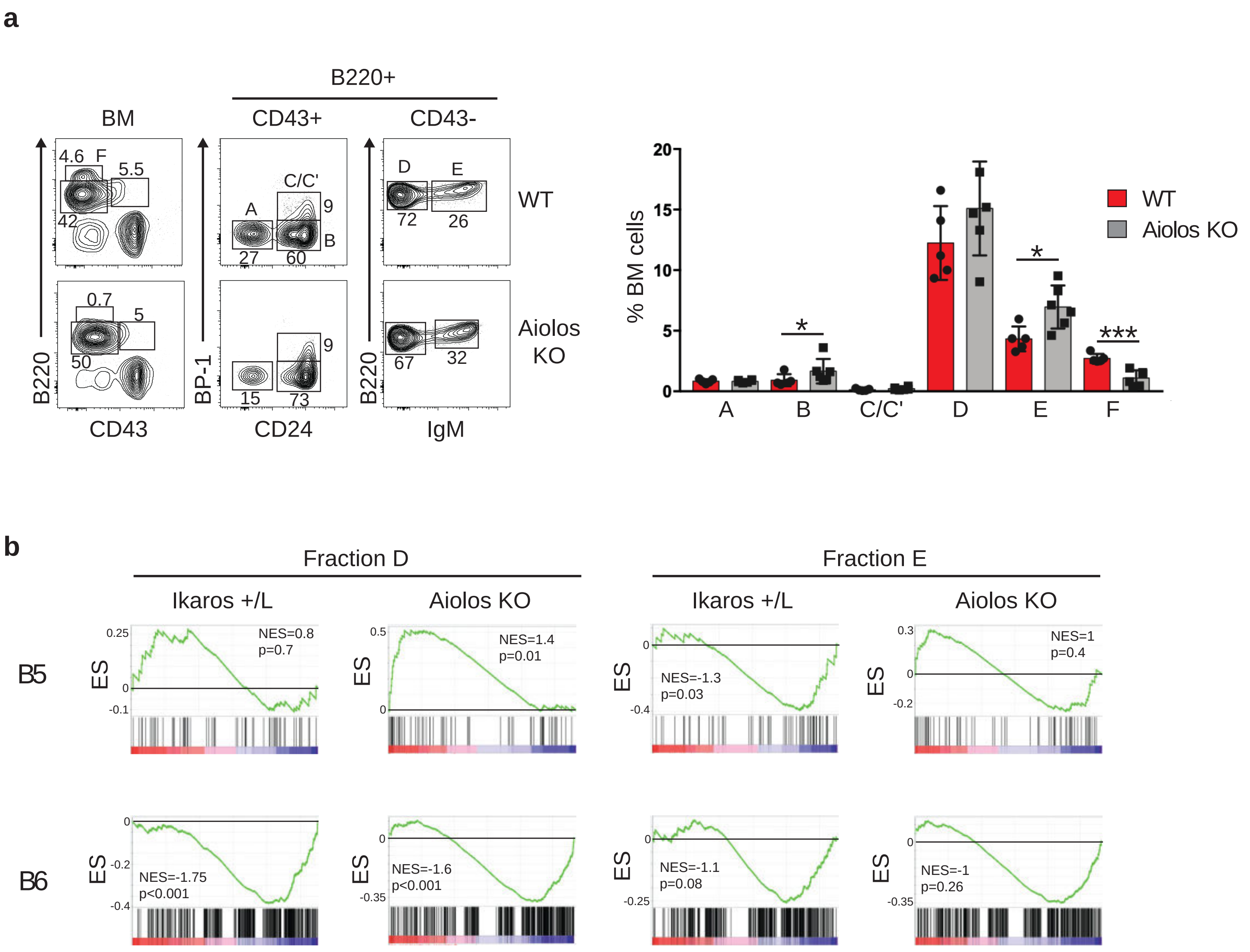


Figure S10
Deau et al

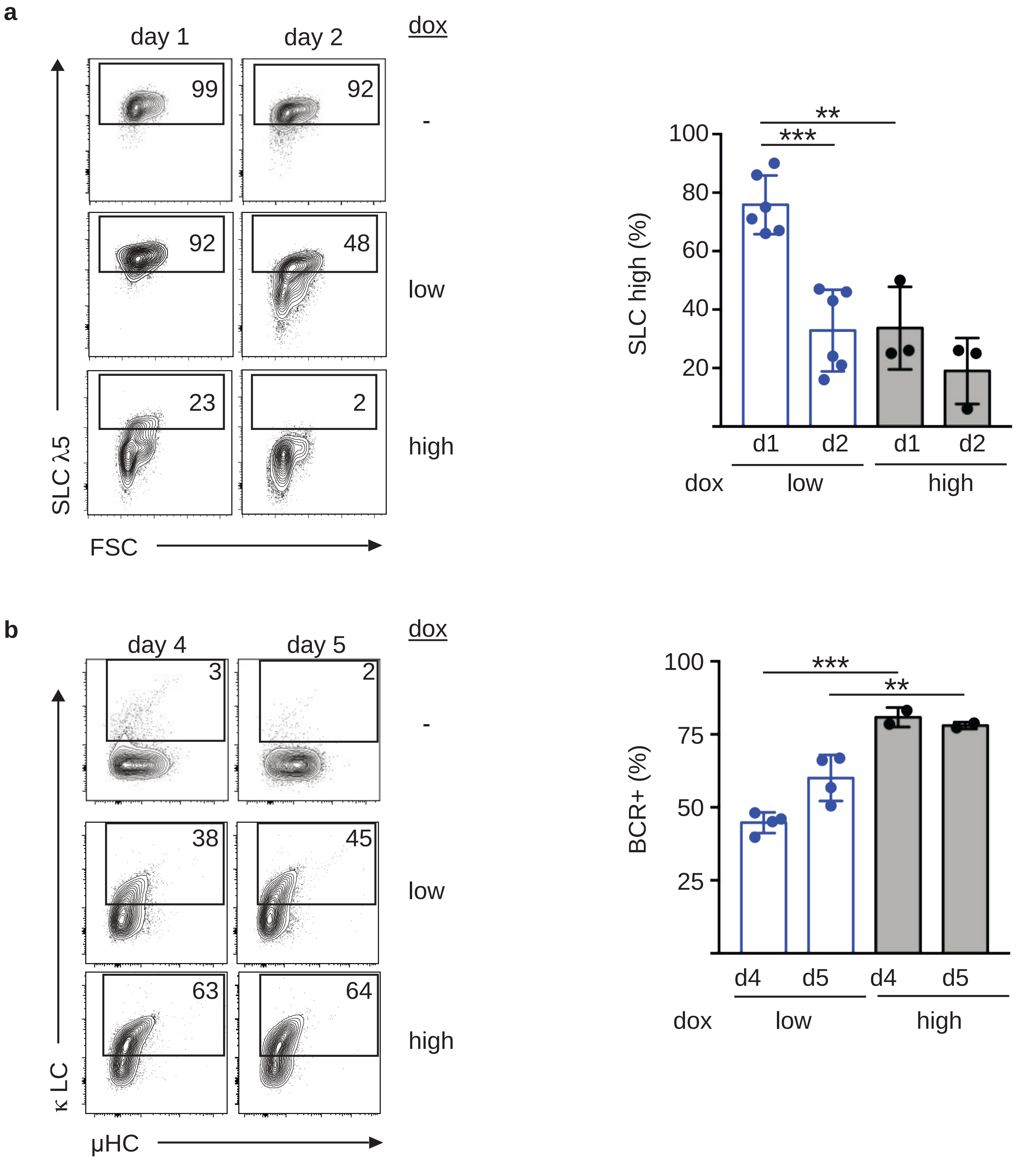


Figure S11
Deau et al
